# Alternative splicing modulation by G-quadruplexes

**DOI:** 10.1101/700575

**Authors:** Ilias Georgakopoulos-Soares, Guillermo E. Parada, Hei Yuen Wong, Eric A. Miska, Chun Kit Kwok, Martin Hemberg

## Abstract

Alternative splicing is central to metazoan gene regulation but the regulatory mechanisms involved are only partially understood. Here, we show that G-quadruplex (G4) motifs are enriched ~3-fold both upstream and downstream of splice junctions. Analysis of *in vitro* G4-seq data corroborates their formation potential. G4s display the highest enrichment at weaker splice sites, which are frequently involved in alternative splicing events. The importance of G4s in RNA as supposed to DNA is emphasized by a higher enrichment for the non-template strand. To explore if G4s are involved in dynamic alternative splicing responses, we analyzed RNA-seq data from mouse and human neuronal cells treated with potassium chloride. We find that G4s are enriched at exons which were skipped following potassium ion treatment. We validate the formation of stable G4s for three candidate splice sites by circular dichroism spectroscopy, UV-melting and fluorescence measurements. Finally, we explore G4 motifs across eleven representative species, and we observe that strong enrichment at splice sites is restricted to mammals and birds.

---

In eukaryotes pre-mRNA processing is key to gene regulation and the generation of isoform diversity. Alternative splicing is arguably the most pivotal mRNA processing mechanism in higher eukaryotes, and in humans it contributes substantially to the protein diversity, affecting 95% of mRNA transcripts (Pan et al. 2008), (Wang et al. 2008), (Licatalosi & Darnell 2010; Oesterreich et al. 2016). Moreover, alternative splicing is essential for normal cell growth, cell death, differentiation, development, sex, circadian rhythms, responses to environmental changes and pathogen responses (Bell et al. 1988), (Kalsotra & Cooper 2011), (Irimia & Blencowe 2012).

Accuracy of pre-mRNA splicing relies on the recognition of three core signals; the 5’ splice site (5’ss), the 3’ splice site (3’ss), and the branch point. Despite the high fidelity observed during the splicing process, computational analyses have reported that human splice site core signals contain only half of the information required to accurately define exon/intron boundaries, implying the involvement of additional sequence features in splice site selection (Lim & Burge 2001), (Wang & Burge 2008). Some of the additional information necessary for splice site definition is found in a complex combination of *cis*-regulatory elements. These splice regulatory elements are short nucleotide sequences which are often bound by RNA-binding proteins (RBPs) that can either facilitate or inhibit the splice site recognition. The role of RBPs and splicing enhancers has been extensively studied, and the current understanding goes a long way towards a quantitative, predictive model of alternative splicing (Barash et al. 2010), (Vuong et al. 2016).

In addition to RBPs, secondary and tertiary structures at both the DNA and RNA level are known to modulate alternative splicing (Shepard & Hertel 2008). More than 20 non-canonical secondary structures have been previously reported for DNA (Ghosh & Bansal 2003), including G-quadruplexes (G4s), hairpins, cruciforms and triplexes. Sequences that predispose the DNA to non-canonical conformations are known as non-B DNA motifs, and they have been characterized with respect to their roles in gene regulation. It has been demonstrated that non-B DNA motifs can influence several aspects, including transcription initiation, transcription termination, and translation initiation (Kumari et al. 2007), (Tellam et al. 2008), (Bugaut & Balasubramanian 2012), (Lam et al. 2013), (Rhodes & Lipps 2015), (Kaushik et al. 2016). Among the non-canonical secondary structures, G4s are the most widely studied class as they have been reported to have an important role in transcriptional regulation of clinically relevant genes. For example, a G4 in the promoter of the oncogene *MYC* acts as a repressor (Siddiqui-Jain et al. 2002), (Hurley et al. 2006), (Yang & Hurley 2006). Similarly, a G4 in the promoter of the proto-oncogene *KRAS* has a negative effect on expression levels (Cogoi & Xodo 2006). Moreover, G4s are also implicated in genomic instability in cancer and neurodegenerative diseases (Siddiqui-Jain et al. 2002), (Wu & Brosh 2010), (De & Michor 2011), (Simone et al. 2015), (Georgakopoulos-Soares et al. 2018).

Many non-B DNA motifs will result in similar secondary structures at the RNA level (Bevilacqua et al. 2016), (Kwok & Merrick 2017), (Strobel et al. 2018). In particular, abundant G4 structure formation in the transcriptome has been demonstrated recently (Kwok et al. 2016). Importantly, the impact of RNA secondary structures in alternative splicing remains only partially understood (Buratti & Baralle 2004), (Warf & Berglund 2010) and although a role of G4s in splicing has been suggested (Hastings & Krainer 2001), (Gomez et al. 2004), (Marcel et al. 2011), (Huang et al. 2017), (Weldon et al. 2018), (Zhang et al. 2019), the extent of G4 impact on alternative splicing remains to be explored. Here we provide the first genome-wide characterization across multiple species of the role of non-B DNA motifs in alternative splicing.

## Results

### Sequence analysis and experimental data show that G4s are enriched near splice sites

To investigate the contribution of non-canonical secondary structures to splice site definition, we systematically explored the distribution of seven known non-B DNA motifs. Since the secondary structures can form both at the DNA and the RNA level (Biffi et al. 2013), (Bevilacqua et al. 2016), (Strobel et al. 2018), we initially considered both strands. These motifs can be identified from the primary sequence, and we focused on the regions flanking human splice sites (Methods). The enrichment profiles varied substantially across the different non-B DNA motif categories (Fig 1a), with exon-intron junctions displaying an acute enrichment for G4s, short tandem repeats and H-DNA motifs. The high enrichment of short tandem repeats was expected since a subset of them overlap with intronic polypyrimidine tracts which are known to be part of the core splicing signal (Dominski & Kole 1991), (Coolidge et al. 1997). By contrast, the enrichment patterns for G4s or H-DNA motifs cannot be explained by the distribution of known splicing signals.

**Figure 1:**
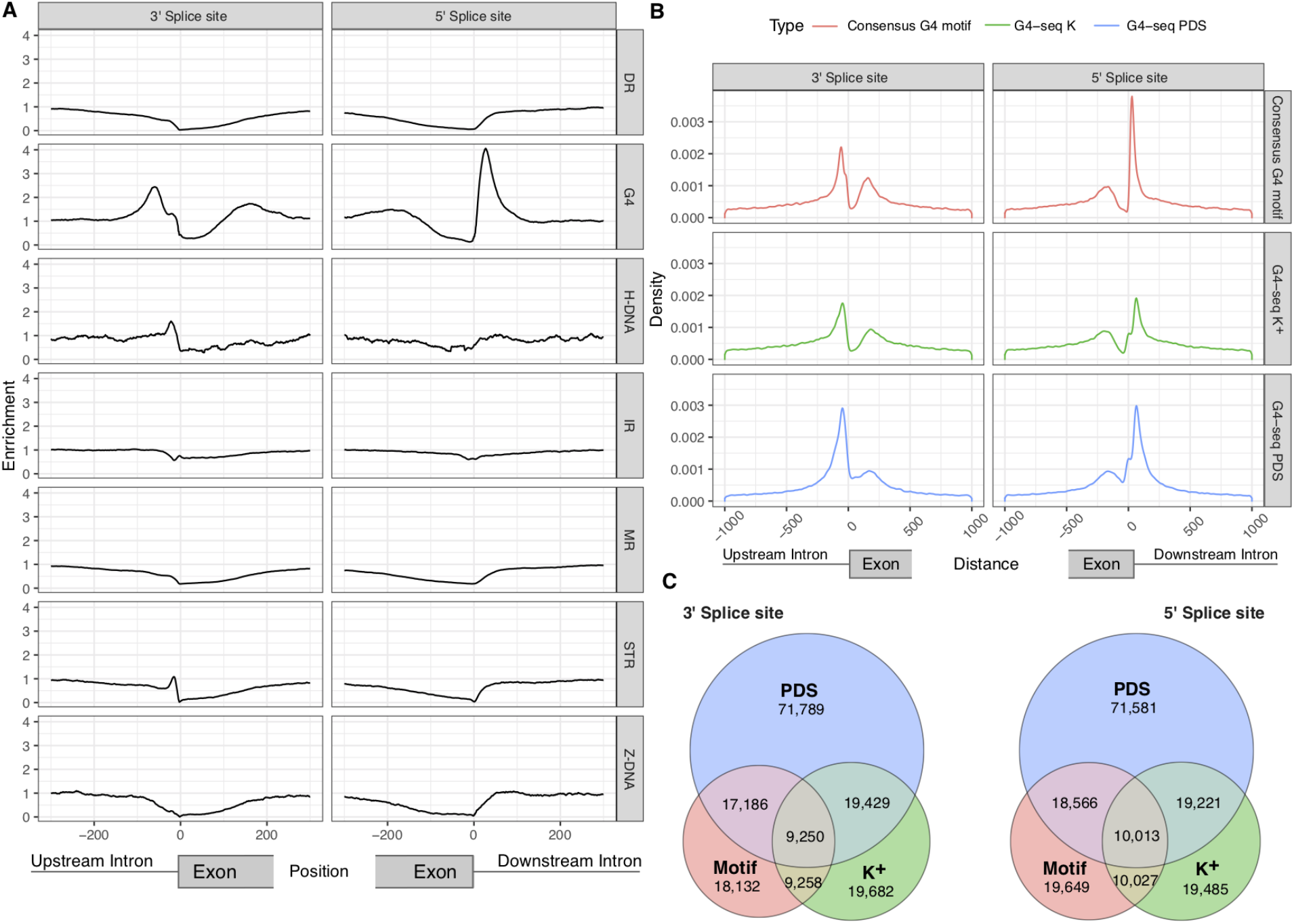
Non-B DNA motifs at splicing junctions. **A.** Distribution of non-B DNA motifs relative to splice sites. Seven non-B DNA motifs are shown, namely direct repeats (DRs), G-quadruplexes (G4s), H-DNA, inverted repeats (IRs), mirror repeats (MRs), short tandem repeats (STRs) and Z-DNA. **B.** Distance between nearest G4 motif / G4-seq peak and a splice site separately for 3’ / 5’ splice sites. **C.** Venn diagrams for the occurrences of G4s within 100 nt of the 3’ss (upstream) and 5’ss (downstream) using the consensus G4 motif, the K^+^ treatment G4-seq derived G4 peaks and the PDS treatment G4-seq derived G4 peaks (Marsico et al. 2019) and reporting the overlapping G4s between them.

The highest enrichment was for G4s, both at the 3’ss (2.44-fold) and the 5’ss (4.06-fold), and this prompted us to further investigate if they have a role in the regulation of splicing. To control for the effect of the nucleotide composition of splice sites in the distribution of the GC-rich G4s, we shuffled the 100 nt window each side of the splice site while controlling for dinucleotide content. Comparing the observed frequency to the median from 1,000 permutations we observed a corrected 2.53-fold and 2.73-fold enrichment for the frequency of G4s at the 3’ss and 5’ss, respectively (p-value<0.001 in both 3’ss and 5’ss), indicating that the G4 patterns are not driven by the sequence composition of splice sites. Moreover, the enrichment was consistent between human and mouse splice sites (Fig 1a, S1a), and the colocalization of G4s and splice sites is not driven by a small number of loci. Within 100 nt of each splice junction we identified 19,987 and 20,088 G4s at the 3’ss and 5’ss, respectively. In total, 31 % of human genes contain a G4 near at least one splice site within a distance of 100 bp. G4s were found within 100 nt for 8.79% and 8.83% of the 3’ss and 5’ss, respectively. The reported G4 frequencies are likely a conservative estimate since we do not take into account intermolecular G4s or G4s that do not adhere to the consensus motif (G⩾3Nl-7G⩾3Nl-7G⩾3Nl-7G⩾3), (Kikin et al. 2006), (Huppert & Balasubramanian 2007), (Varizhuk et al. 2017).

To evaluate if the G4 motifs that were enriched near splice sites lead to the formation of DNA G4 structures *in vitro,* we analyzed previously published G4-seq data (Chambers et al. 2015), (Marsico et al. 2019). G4-seq utilizes the fact that stable G4s can stall the DNA polymerase *in vitro,* thereby allowing high throughput sequencing to be used to detect G4s at high resolution. We first measured the distribution of G4s relative to the splice sites for HEK-293T cells in Pyridostatin (PDS) and K^+^ treatments from (Marsico et al. 2019). In both conditions, we observed an enrichment of G4-seq peaks relative to the 3’ss and 5’ss, but with a more pronounced G4 enrichment in PDS treatment compared to K^+^ treatment (Fig 1b). The majority of G4 positions derived from G4-seq peaks in K^+^ and PDS treatments did not overlap consensus G4 motifs (Fig 1c). Since the G4-seq assay can also identify G4s with non-canonical motifs, it is to be expected that the overlap with the consensus G4 motifs would be limited. We replicated our results in primary human B lymphocytes (NA18507) in Na^+^-K^+^ and Na^+^-PDS conditions (Chambers et al. 2015), both of which promote G4 formation. In both conditions, only ~25% of G4-seq derived peaks are captured by the consensus G4 motif. Nevertheless, at splice sites we found an enrichment comparable to that obtained from the motif analysis, directly implicating G4 formation at splice sites (Fig S1b-d). Differences between the G4-seq datasets are likely the result of the differences in the experimental settings and treatments between the two studies (Chambers et al. 2015), (Marsico et al. 2019). For both the PDS and K^+^ treatments we find that a substantial fraction of the genome is affected, with 31.72% and 10.25% of splice junctions having a G4 nearby. In addition, 67% and 35% of human genes contain a G4-seq peak from PDS and K^+^ treatments within 100 nt of a splice junction, supporting our earlier observations using the consensus G4 motif. As a result of these findings, we conclude that G4s are a pervasive feature across splicing junctions.

### G4s are preferentially found on the non-template strand

Since G4s are strand specific, we oriented each instance relative to the direction of transcription. Thus, we considered G4s found at the template (non-coding) and non-template (coding) strands separately and found them statistically enriched on the non-template strand (Binomial tests, p-value<0.001 at 3’ss and 5’ss). G4s were enriched at both strands. At 3’ss the enrichment was 3.01-fold and 2.78-fold enrichment scores at the non-template and template strands, respectively (Fig S2). At the 5’ss the difference between the strands was larger with 5.56-fold and 2.38-fold at the non-template and template strands, respectively (Fig S2). Therefore, there was an asymmetric enrichment between the template and non-template strands at the 5’ss, but only a weak asymmetry at the 3’ss.

### G4s are enriched at weak splice sites

Weak splice sites are highly involved in alternative splicing and often contain additional regulatory elements (Parada et al. 2014), (Sibley et al. 2016), (Erkelenz et al. 2018). To explore the distribution of G4s across weak and strong splice sites, we calculated a splicing strength score for all internal exons based on splice site position weight matrices (Sheth et al. 2006), (Parada et al. 2014). We grouped splice sites into four quantiles based on the splicing strength scores, and explored the enrichment levels of G4 motifs for each quantile separately. We found an inverse relationship between the calculated splicing strength score and G4 enrichment, with the weakest splice sites having the highest enrichment of G4s both at the 3’ss and the 5’ss with 2.77-fold and 4.95-fold enrichment, respectively (Fig S3). For both mouse and human, the splicing strength scores for splice junctions with a G4 are significantly lower than for splice junctions without a G4 (Mann-Whitney U, p-value<0.001). The same inverse relationship between the splicing strength score at splice sites and the G4 enrichment is found in the G4-seq data for human in PDS and K^+^ treatment (Marsico et al. 2019) and for PDS-Na^+^ and K^+^-Na^+^ treatment (Chambers et al. 2015), (Fig S4a-d), (Mann-Whitney U, p-values<0.001).

We also investigated if there was a strand asymmetry when considering the splicing strength scores. Indeed, we found a bias in the splicing strength scores dependent on the strand orientation of G4s (Mann-Whitney U, 3’ss p-value<0.05, 5’ss p-value<0.001). At the 3’ss the enrichment for splice junctions with the weakest splicing strength scores at the template and non-template strand was 3.90-fold and 3.66-fold, respectively. By contrast, we observed a 6.76-fold enrichment for G4s at the 5’ss at the non-template strand, but only a 3.66-fold enrichment on the template strand at the splice junctions with the weakest splicing strength scores (Fig 2a). Taken together, these results indicate a preference for the non-template strand that is inversely proportional to the splicing strength score (Fig 2a). They also suggest that G4s are more prevalent at the non-template strand, which is the sequence found in the transcribed mRNA. We validated the differences in the distribution of G4 motifs at the template and non-template strands and the associated differences in splicing strength score using G4-seq from the two available datasets (Chambers et al. 2015), (Marsico et al. 2019). Consistent with the results derived from the consensus G4 motif, we found a preference for the nontemplate strand at the 5’ss of weak splice sites using the G4-seq datasets (S4e-h).

**Figure 2:**
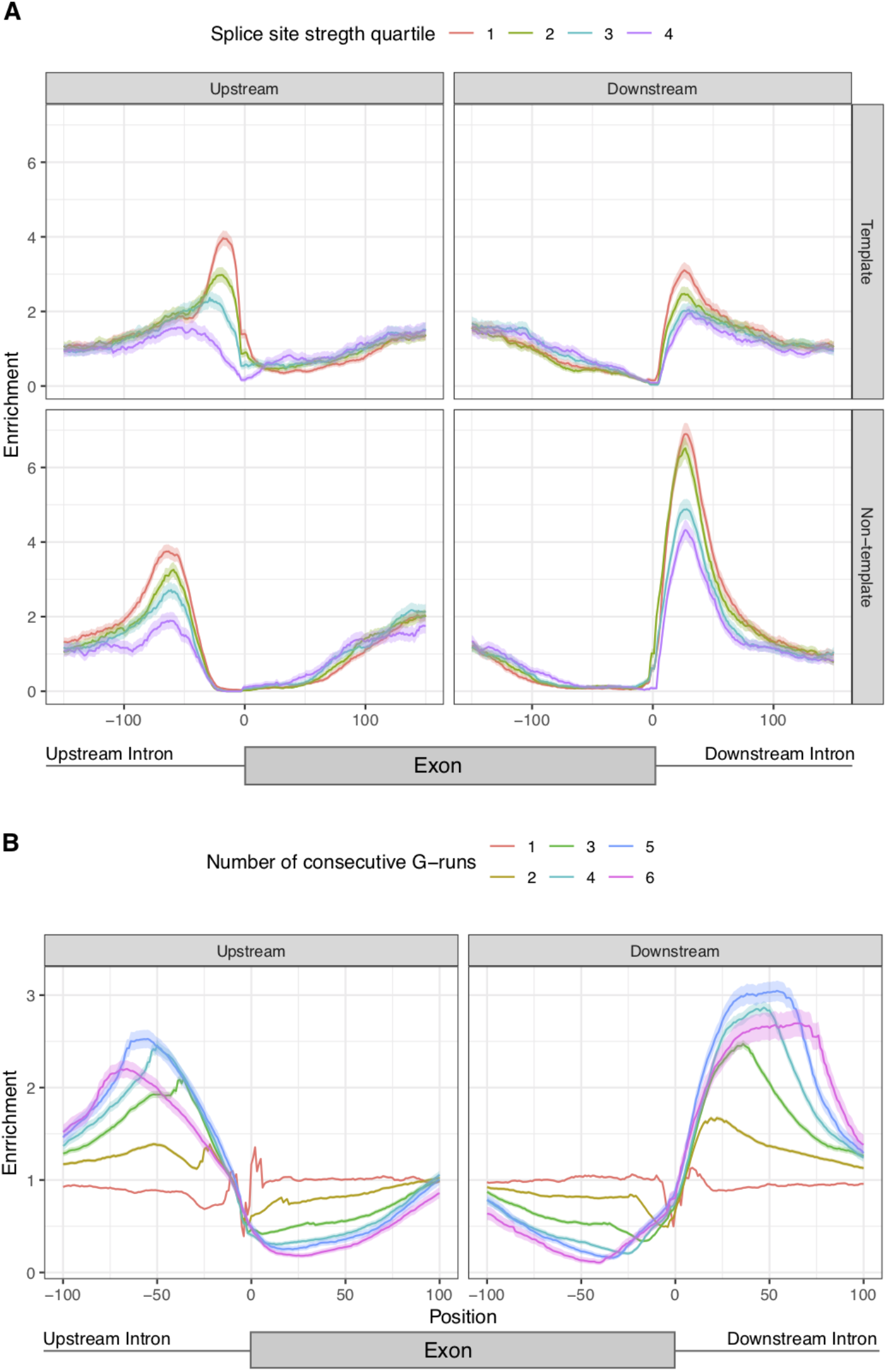
Characterisation of G4 positioning across splicing junctions. **A.** G4 enrichment for template and non-template strands and stratified by the splicing strength scores of the adjacent splice site. The splicing strength scores for splice junctions with a G4 are significantly lower than for splice junctions without a G4 (Mann-Whitney U, p-value<0.001). The splicing strength score bias was found to be dependent on the strand orientation of G4s for splice sites with G4s within 100 nt away (Mann-Whitney U, 3’ss p-value<0.05, 5’ss p-value<0.001). **B.** Number of consecutive G-runs and relative enrichment at the splicing junction. The error bands in A-B represent 0.95 confidence intervals from the binomial error.

### Longer G-runs are more highly enriched at splice sites

An intramolecular G4 is usually a representation of four or more consecutive G-runs. Yet, intermolecular G4s can form with fewer G-runs since multiple molecules can contribute to G4 structure formation (Bhattacharyya et al. 2016). We found minimal to no enrichment for single G-runs at both 5’ss and 3’ss (Fig 2b). However, for two and three G-runs we observed a 1.39-fold and a 2.10-fold enrichment at the 3’ss and a 1.67-fold and a 2.47-fold enrichment at the 5’ss, which may implicate intermolecular G4s in splice sites. The highest enrichment was observed for four to six G-runs, indicating that intramolecular G4 motifs are more enriched at splice sites than their intermolecular counterparts, in accordance with our earlier findings.

### G4s are enriched for short introns

The length of introns in metazoans can vary across four orders of magnitude (Sakharkar et al. 2004). We hypothesized that the enrichment patterns of G4s at introns proximal to splicing sites would be associated with intron length. We compared the intron length of splice sites that had a G4 within 100 bps in the direction of the intron to the ones that did not have this motif. Consistent with our hypothesis, we found that introns with a G4 at the 3’ss had a median length of 701 nt while introns without a G4 had a median length of 1,618 nt (S5a), (Mann Whitney U, p-value<0.001). Similarly, at the 5’ss, introns with a G4 had a median length of 379 nt, whereas introns without a G4 had a median length of 1,629 nt (Mann Whitney U, p-value<0.001). Interestingly, introns in the range of ~45-85 bps were the most enriched for G4s for both the 3’ss and the 5’ss. Moreover, the enrichment of introns in G4s declined rapidly with increased intron length, indicating that they are preferentially found in the subset of short introns (Fig 3a-b, Kolmogorov-Smirnov test p-value<0.001). We also investigated the association between splicing strength score and intron length at sites with G4s in the 3’ss and 5’ss and found that the highest enrichment for G4s was in short introns with weak splice site strength (Fig 3c). However, when comparing G4 enrichment at splice sites of short introns with a selection of long introns that have the same GC-content distribution, we found no difference or even higher enrichment of G4 at long intron splice sites (Fig 3d, S5b), indicating that the association of G4 to splice sites located a short intron could be driven by GC-content.

**Figure 3:**
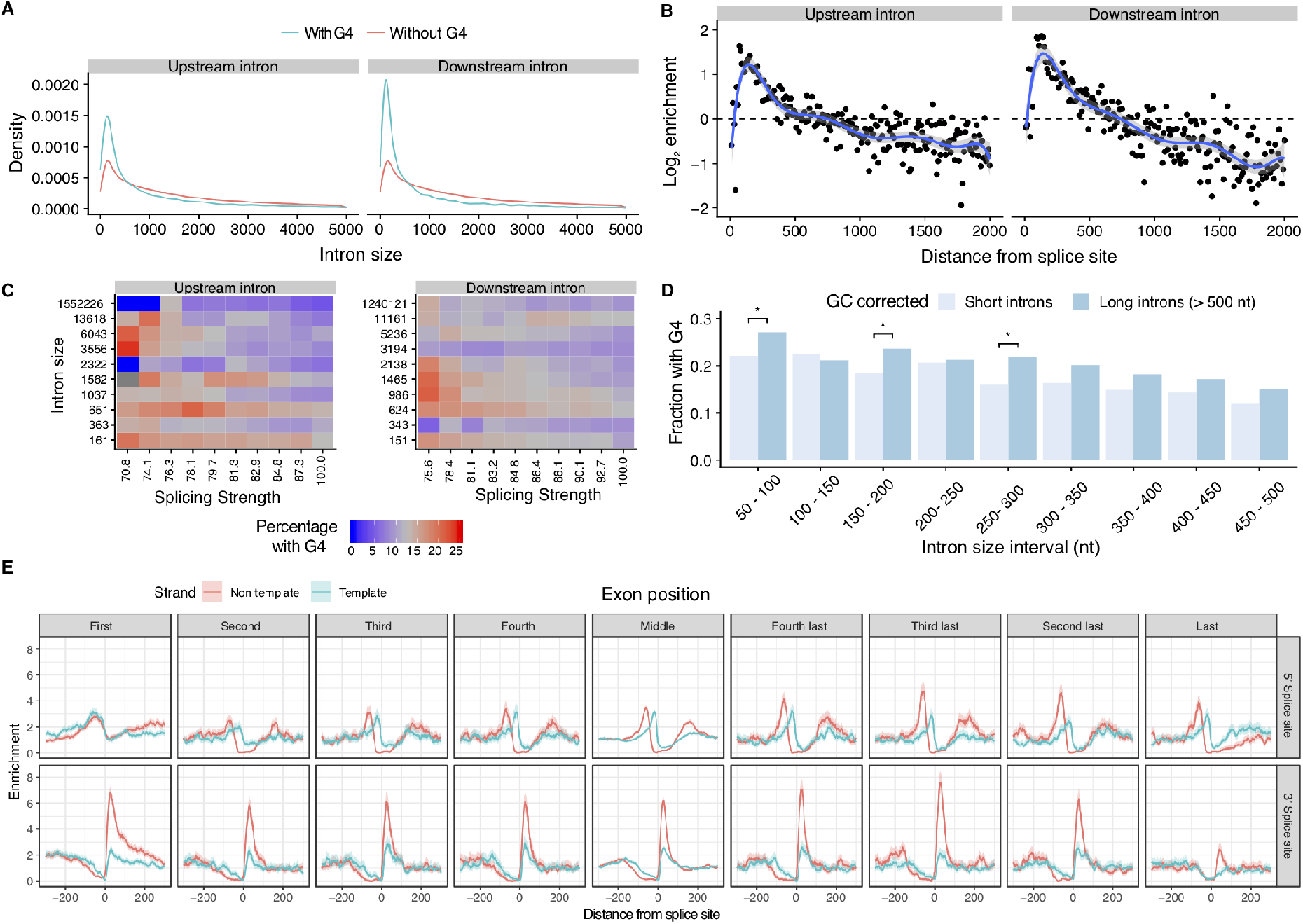
Distribution of G4s in the vicinity of splicing sites. **A.** Length density distribution of introns upstream and downstream from exons that are flanked or not by G4s (Kolmogorov-Smirnov test p-value<0.001). **B.** Abundance enrichment of intron sizes at upstream and downstream splice sites flanked by G4s. A bin size of 10 bps was used with the blue line representing an eighth-degree polynomial model. **C.** Heatmap for the relationship between splicing strength score, intron length and G4 presence in a local window of 100 nt within the splice site for the upstream and downstream introns. Red color represents high proportion of splice site regions with G4s, whereas blue color represents depletion of G4s. **D**. Fraction of splice sites with a G4, controlling for GC content between long and short introns. We use Chi-squared test to evaluate significant differences between short and long introns (* denotes p-values<0.05 after multiple testing correction). **E.** G4 motif enrichment relative to the splice site across exons in the gene body at the 3’ss and at the 5’ss for template and non-template strands. G4 motifs are enriched at both 3’ss and 5’ss across splice sites throughout the gene body. Exons were separated into first to fourth exons, middle exons, last four exons and the distribution of G4s was studied individually for each category.

To further investigate the relationship regarding the intron length, we separated G4s identified using the consensus motif into non-template and template for both the 5’ss and the 3’ss. In this case, the GC-content is not a covariate since the template and non-template strands have the same GC-content. At the 3’ss introns showed small but significant differences in length if a G4 was at the non-template or the template strand with medians of 736 nt and 621 nt, respectively (Mann-Whitney U, p-value<e-21). However, if a G4 was at the non-template strand at the 5’ss the median intron length was 267 bp, whereas if the G4 was at the template strand the median intron length was 539 nt (Mann-Whitney U, p-value=e-288), displaying more aggravated differences in intron length. Therefore, we conclude that the highest enrichment is found for short introns, on the non-template strand, downstream of the 5’ss.

We also investigated if there is an association between G4s near splice sites and exon length. We do not find a significant association between G4s and exon length at the 3’ss (median exon length without G4s: 124 bp, median exon length with G4s: 123 bp, p-value>0.05, Mann-Whitney U), but we find a significant association for smaller exons near the 5’ss, albeit with a very small magnitude (median exon length without G4s: 127 bp, median exon length with G4: 123 bp, p-value<0.001, Mann-Whitney U). Furthermore, we explored if microexons, defined as exons <30 nt long (Irimia et al. 2014), (Li et al. 2015) had an enrichment for G4s at their splice sites relative to other exons. However, we could not find a higher density of G4s at the introns flanking microexons than other exons.

In addition to exploring the relationship between intron and exon lengths and G4s, we also considered the position across the gene body. For each gene with nine or more exons, we separated the exons of its longest transcript into nine groups; the first four exons, the last four exons and the remaining middle exons. We find an attenuation of the enrichment around the last intron for the 5’ss and at the first couple of introns for the 3’ss, indicating clear differences at both ends of the transcript in comparison to other introns (Fig S5c), this result indicates that the role of G4s in splicing regulation is pervasive across the gene body.

Importantly, we also measured separately the G4 enrichment at the template and nontemplate strands of splice junctions across the gene length (Fig 3e). We found that the enrichment of G4s was consistently higher in the non-template strand across exons. The foremost difference between enrichment score for the two strands was found in the 3’ss exceeding 3-fold higher enrichment, while the differences between the two strands at the 5’ss were smaller. These findings provide evidence for widespread variation in the topography of G4s in splice junctions; these include the frequency of G4s in the exons and introns flanking the splice site, biases regarding the strand preference, the distance from the splice site and the positioning across the gene body.

### Dynamic splicing responses to stimuli are associated with proximal G4s

To gain further insights regarding the biological roles of G4s in the modulation of alternative splicing events, we investigated if G4s are associated with dynamic splicing changes in response to depolarisation stimuli. To that end we analyzed published data from mouse and human embryonic stem cell (ESC)-derived neurons and mouse primary neurons subjected to a depolarisation solution including 170mM of KCl and an L-type Ca2^+^ channel agonist, resulting in an influx of Ca2^+^ and followed by RNA-seq four hours post-treatment (Qiu et al. 2016). The rise of intracellular Ca2^+^ has been shown to have an impact on alternative splicing mediated by calmodulin dependent kinase IV (CaMK IV), (Sharma & Lou 2011).

We used Whippet (Sterne-Weiler et al. 2018) to quantify alternative splicing changes after depolarisation and compared to the unstimulated controls (Methods) (Supplementary Material). The change in inclusion of an exon is quantified using percent spliced included (PSI) which represents the fraction of transcripts that contain the exon. As an illustration we considered exons flanked by one or more G4s in the SLC6A17, UNC13A and NAV2 loci that were differentially included after treatment for further experimental validation (Fig 4a, S6). Firstly, *SLC6A17* (NTT4/XT1) is a member of SLC family of transporters which are involved in Na^+^-dependent uptake of the majority of neurotransmitters at presynaptic neurons (Zaia & Reimer 2009). SLC6A17 is involved in the transport of neutral amino acids and mutations in this gene have been associated with Autosomal-Recessive Intellectual Disability (Zaia & Reimer 2009), (Iqbal et al. 2015). We show that exon seven from SLC6A17, which is alternative skipped after KCl treatment (Delta PSI=-0.177), has a G4 50 nt downstream of the 5’ss on the non-template strand. As the domains of SLC6A17 include an intracellular loop, two transmembrane regions and part of extracellular domains, the KCl-induced alternative skipping of this exon may lead to functional structural changes (Figure 4a). Similarly, *UNC13A* encodes another presynaptic protein involved in glutamatergic transmission and it has been associated with amyotrophic lateral sclerosis (Placek et al. 2019). We identify a G4 downstream of exon 38, which results in dramatic exon skipping (Delta PSI=-0.369), (S6a).

**Figure 4:**
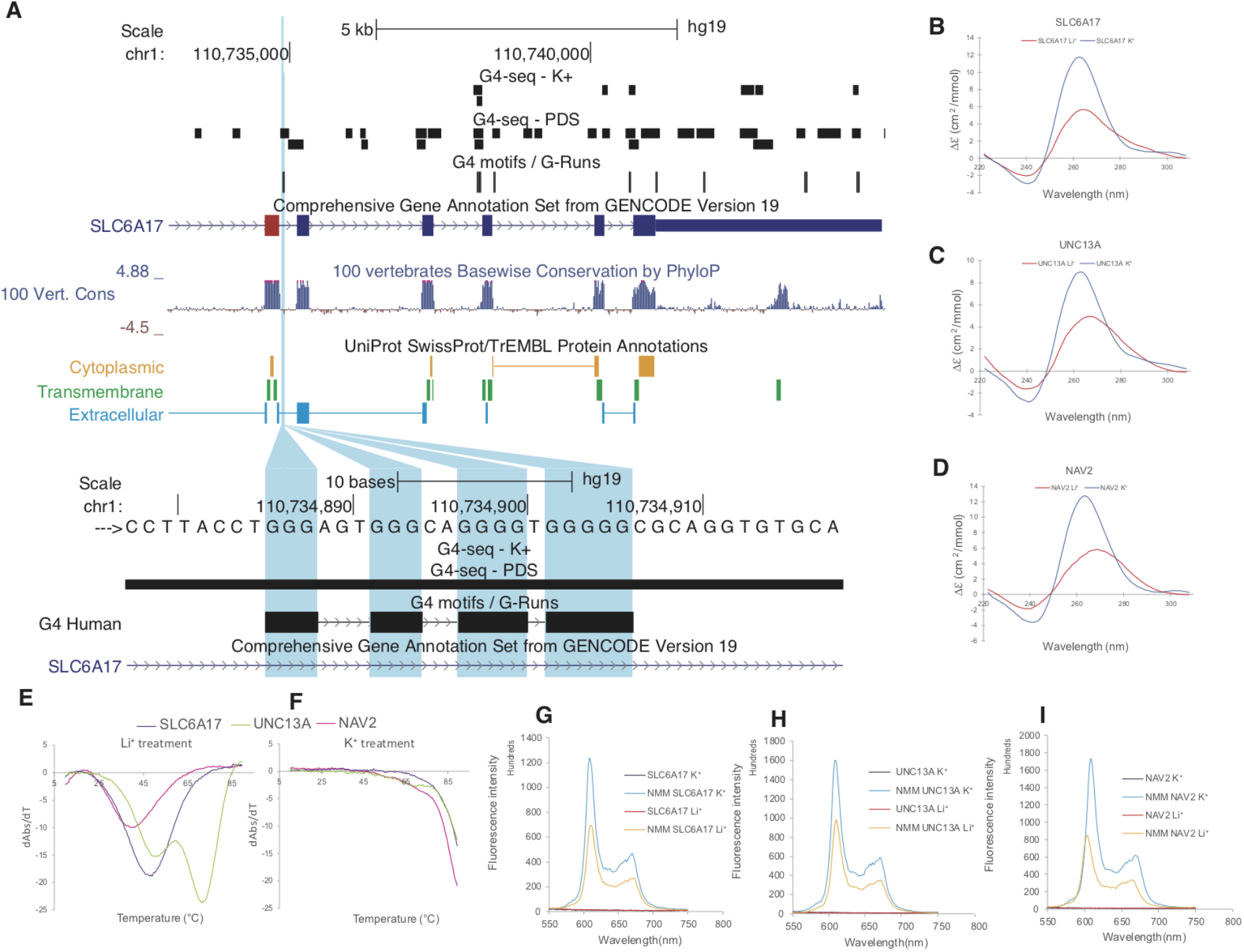
Targeted validation of RNA G4s found in splicing sites in presence of a G4 stabilising cation (K^+^) and a non-G4 stabilising cation (Li^+^). **A.** UCSC screenshot displaying the *SLC6A17* locus and the distribution of G4 consensus motifs, G4-seq derived G4 peaks and protein domains. **B-D.** Circular dichroism (CD) spectra of the three candidate targets for G4 formation potential in presence of two cations. The monovalent ion-dependent nature (G4 stabilized in K^+^ but not in Li^+^) and the CD profile (positive peak at 262nm and negative peak at 240nm) indicate the formation of RNA G4s with parallel topology. **E-F.** UV melting profiles of the three G-quadruplex candidates in presence of Li^+^ and K^+^. Hyperchromic shift at 295nm is a hallmark for G4 formation, which can be transformed to negative peak in derivative plot (dAbs/dT) for G4 stability analysis. The melting temperature (Tm) of a G4 can be identified at the maximum negative value. For the K^+^ treatment, the Tms of the G4 are >85°C. **G-I.** Fluorescence emission associated with NMM ligand binding to G4 candidates in the presence of Li^+^ or K^+^ ions. In the absence of NMM ligand, no fluorescence was observed at ~610 nm. Upon NMM addition, weak fluorescence was observed under Li^+^, which was substantially enhanced when substituted with K^+^, supporting the formation of G4 which allows recognition of NMM and enhances its fluorescence.

Finally, the third target was a G4 located downstream of exon 16 in NAV2 (navigator protein 2), which is required for retinoic acid induced neurite outgrowth in human neuroblastoma cells (Merrill et al. 2002). Again, KCl treatment resulted in exon skipping (Delta PSI=-0.271), which affects a NAV2 serine rich sequence region (S6b).

For each of the three candidates we performed multiple assays to provide additional support for the formation of G4s at the RNA level (Kwok & Merrick 2017), (Chan et al. 2018), (Chan et al. 2019). First, we performed circular dichroism spectroscopy and UV-melting measurements of the G4-containing RNA oligonucleotides, in the presence of lithium ions (non-G4 stabilising) or potassium ions (G4 stabilizing), to examine the formation potential and stability of RNA G4s. Supporting the case that G4s are present in the transcripts, we found that all three candidates folded into stable RNA G4 structures preferentially under K^+^ conditions (Fig 4b-f, S7a). To confirm the results from the circular dichroism and UV-melting experiments, we further used fluorescent-based arrays such as *N*-methyl mesoporphyrin IX (NMM) ligand enhanced fluorescence, Thiovlavin-T (ThT)-enhanced fluorescence, and intrinsic fluorescence experiments (Fig 4g-i, S7b-c). Indeed, we observed increased fluorescence intensity under conditions that promoted G4 formation for all the three candidates, confirming the formation of RNA G4s in these examples.

Having validated the formation of G4s around three exons that are differentially included after KCl induced depolarisation, we examined their impact on splicing genome wide (Fig 5a). We find that exon skipping at core exons is associated with G4s (chi-squared test multiple testing corrected, p-value<5e-12 Fig 5a). We also found inclusion of alternative first exons following KCl treatment indicating alternative promoter usage (chi-squared test, multiple testing corrected, Fig 5a). The analysis identified a total of 22,344 splicing events where the absolute value of Delta PSI was greater or equal than 0.1 and the probability was greater or equal to 0.9. We focused our analysis on the 2,633 events that involved cassette exons, and of these 2,346 (89.1%) corresponded to increased skipping (Fig 5b, binomial test, p-value<0.001). These results are consistent with previous studies which have demonstrated exon skipping following depolarisation in individual examples (An & Grabowski 2007), (Lee et al. 2007), (Schor et al. 2009), (Liu et al. 2012), (Fiszbein & Kornblihtt 2017), but to the best of our knowledge we provide the first genome-wide analysis demonstrating widespread exon skipping (Fig 5b). Interestingly, we found an enrichment for differential splicing events with G4 motifs at the associated splicing sites, exceeding that which was expected by the background distribution (Fig 5a, c, chi-squared test with multiple testing correction, p-value<0.001, odds-ratio=1.57). To provide further support for the findings obtained from the consensus G4 motif, we examined the distribution of G4-seq derived peaks in PDS and K^+^ conditions around splice sites of differentially and non-differentially included exons. As expected, we found a consistent enrichment at the differentially included exons (Fig 5a, d-e). Moreover, the effect size was larger for the G4 motifs and the G4-seq derived G4 sites in the non-template strand at the 5’ss in comparison to those found at the template strand (chi-square test multiple testing correction, p-value<0.001 when using the consensus G4 motif and for both PDS and K^+^ G4-seq conditions in human neurons). In addition to human cells, we performed the same analysis in similar experiments in mice across four different conditions (Fig S8–S11). We recapitulate the widespread exon skipping phenomenon observed after depolarisation in human neuronal cells (Fig 5a, S8), but we found a significant association of alternative included exons only with PDS G4-seq derived peaks in two conditions (Fig S9–S11). Importantly, we report alternative promoter usage associated with G4s and alternative first exon inclusion in both mouse and human neurons following KCl treatment (Fig 5a, S9). Taken together, our results suggest that the presence of a G4s at the splice junction of cassette exons is associated with dynamic changes of alternative splicing in response to KCl induced depolarisation.

**Figure 5:**
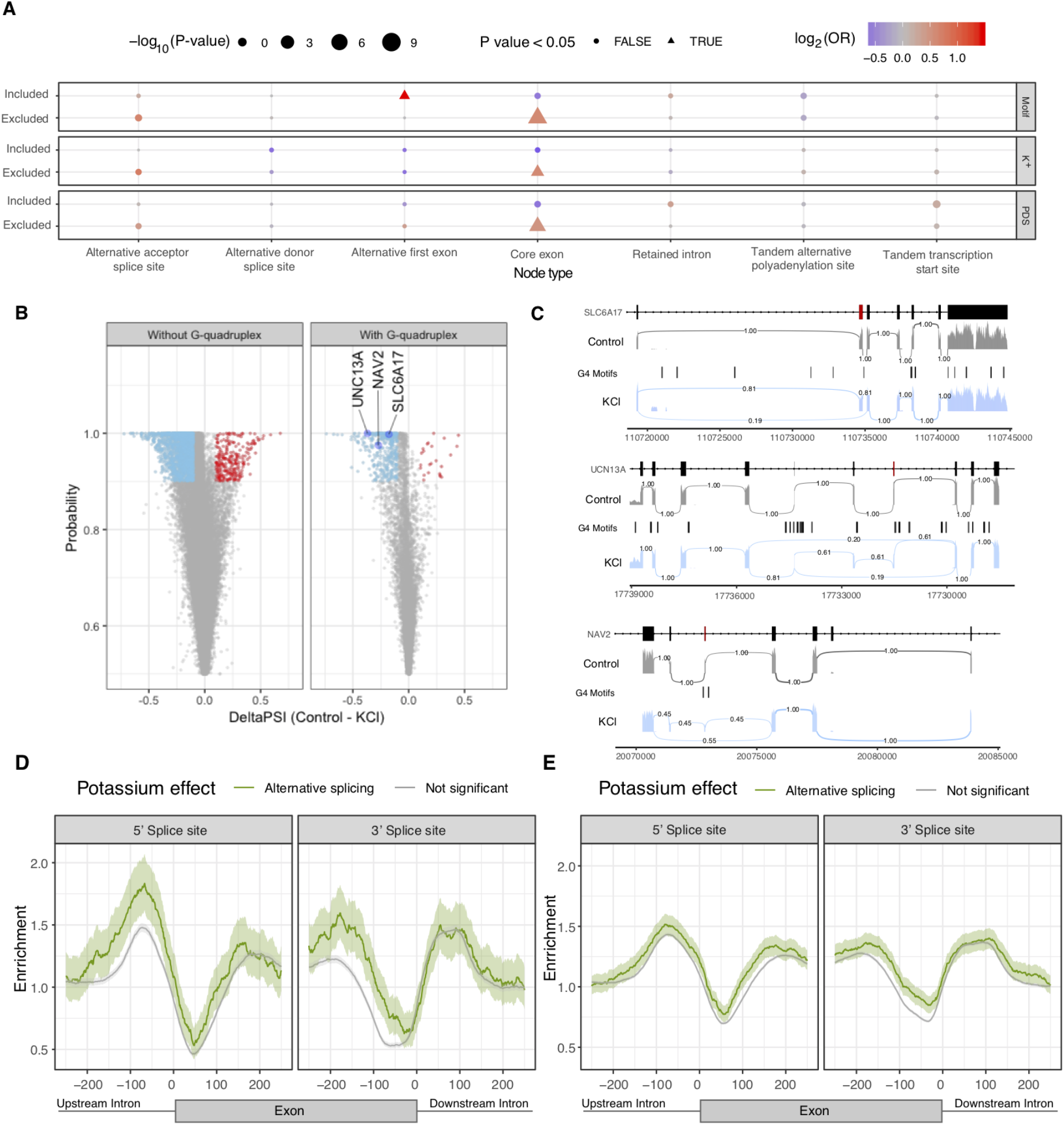
Neuronal stimulation with KCl results in alternative splicing events associated with presence of G4s. **A.** G4 association with alternative splicing changes. Inclusion and exclusion patterns of splice nodes are shown in association to G4 presence or absence following KCl treatment. Odds ratio representing the relationship between presence of G4s and alternative splicing changes. The odds-ratio significance was assessed by chi-squared tests. All p-values were calculated with chi-squared tests using Yates’ correction and also adjusting for multiple testing with Bonferroni corrections. **B.** Volcano plot showing differential inclusion events in presence and absence of flanking G4s and the associated probability following potassium stimulation with widespread exon skipping after depolarisation in human neuronal cells. **C.** Sashimi plots showing alternative exon inclusion for the three candidates, namely SLC6A17, UNC13A and NAV2 following KCl treatment. Exons flanked by a G4 that were used for validation experiments (Fig 4, S6) are marked in red. The numbers connecting exons represent the fraction of reads supporting each path. **D.** Enrichment plots at 3’ / 5’ splicesite vicinity after potassium stimulation of human neurons using G4-seq derived G4 maps with PDS treatment. Splice sites with G4 peaks within 100 nt were more likely to be differentially spliced following KCl treatment (Chi-squared test, p-value<0.001 both at 5’ss and 3’ss). **E.** Enrichment plots at 3’ / 5’ splice site vicinity after potassium stimulation of human neurons using G4-seq derived G4 maps with K^+^ treatment. Splice sites with G4 peaks within 100 nt were more likely to be differentially spliced following KCl treatment (Chi-squared test, p-value<0.001 both at 5’ss and 3’ss). The error bands in D-E represent 95% confidence intervals based on a binomial model.

### Enrichment of G4s at splice sites does not extend beyond vertebrates

Alternative splicing is a pivotal step of eukaryotic mRNA processing. To understand to what extent splice site regulation by G4s is conserved we considered eleven eukaryotes: *Homo sapiens* (human), *Mus musculus* (mouse), *Sus scrofa* (pig), *Gallus gallus* (chicken), *Danio rerio* (zebrafish), *Caenorhabditis elegans* (nematode) *D. melanogaster* (fruit fly), *Xenopus tropicalis* (frog), *Anolis carolinensis* (lizard), *Saccharomyces cerevisiae* (yeast) and *Arabidopsis thaliana* (flowering plant). *S. cerevisiae* was excluded from further analysis since we could not find any G4s at splice sites and G4s were rare with only 39 occurrences genomewide. Interestingly, we found that the enrichment pattern of G4s at splice sites was restricted to a subset of vertebrate species, with minimal or no enrichment in fruit fly, *Arabidopsis* and *C. elegans* (Fig 6a-b, S12a). We observed strong enrichment in chicken, pig, human and mouse, while lizard displayed limited enrichment levels. Surprisingly, frog and zebrafish displayed relative depletion. This suggests that alternative splicing regulation by G4s is found is restricted to mammals and birds, but absent in plants, other tetrapods or fish.

**Figure 6:**
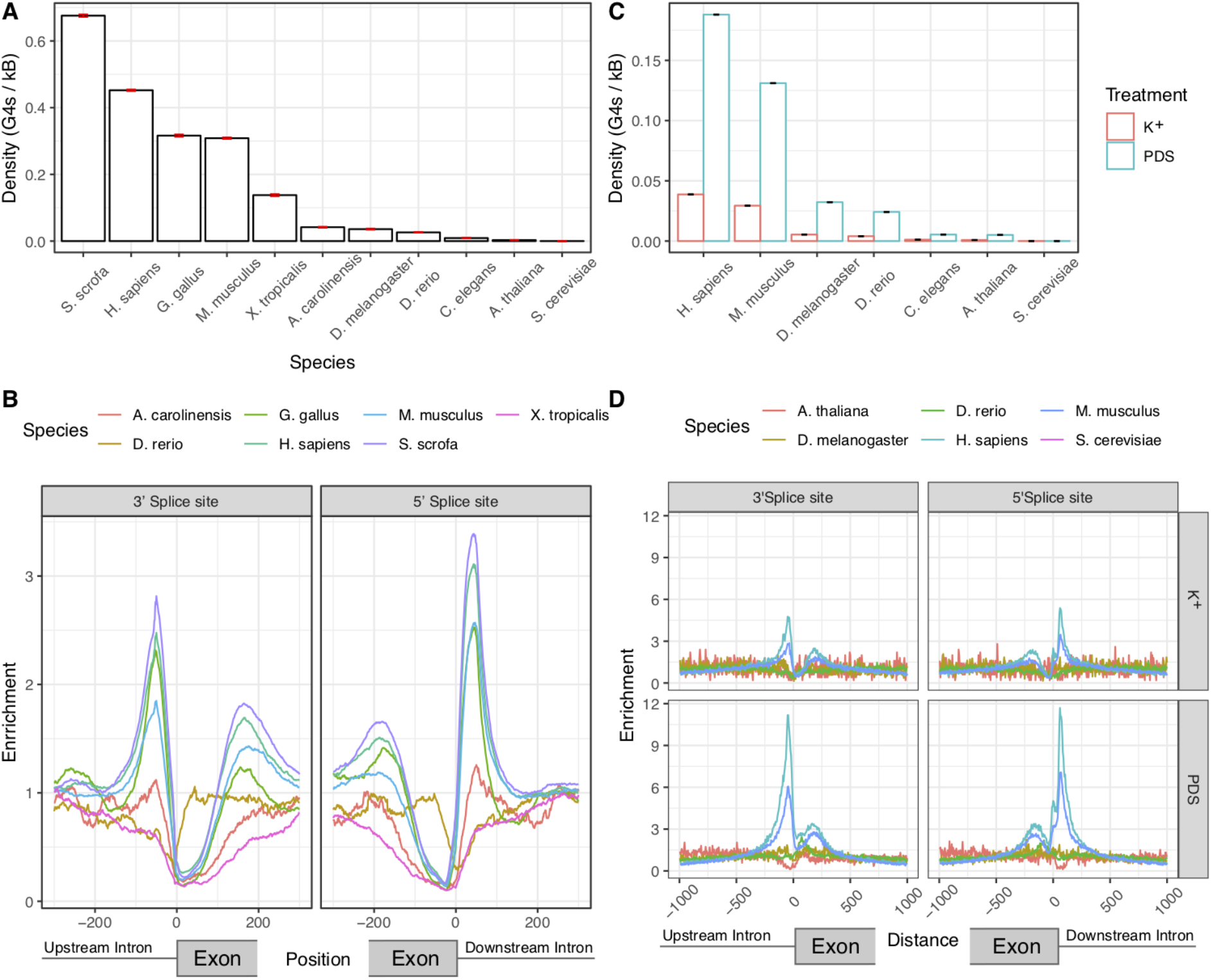
Enrichment of G4s at splicing sites across different species. **A.** Density of G4 motifs in a 100 nt window each side across all 5’ / 3’ splice sites of each species. Error bars indicate standard deviation from 1,000-fold bootstrapping with replacement. **B.** Enrichment of G4 motifs at splice sites for seven vertebrate species, using the consensus G4 motif. **C.** Enrichment of G4-seq derived G4s at splicing sites at 100 nt splicing site windows in PDS and K^+^ treatments. Error bars indicate standard deviation from 1,000-fold bootstrapping with replacement. **D.** Enrichment of G4s at: 5’ / 3’ splice sites across six species for PDS and K^+^ treatments.

Additional support for this conclusion comes from our analysis of G4-seq derived G4 maps generated in PDS and K^+^ conditions. These maps are available for multiple model organisms, including three vertebrates (human, mouse and zebrafish) and four non-vertebrate species (nematode, fruit fly, arabidopsis and yeast). Consistent with the analysis based on the primary sequence, we find an acute enrichment of G4s at the 5’ss and 3’ss only in humans and mouse. In particular, we could not find any G4s in the vicinity of splicing junctions for *S. cerevisiae,* there was no enrichment for *D. melanogaster* and *D. rerio,* while we observed a depletion in *A. thaliana* (Fig 6c-d, S12b).

## Discussion

The identification of regulators of splicing remains an active area of research, as the information content at splice sites is insufficient for predicting alternative splicing events in higher eukaryotes (Lim & Burge 2001), (Barash et al. 2010). Here, we provide evidence for a widespread role of G4s in the regulation of splicing. The enrichment of G4s at splice junctions is comparable to what is observed at promoters in human (Fig S12c), even though it is primarily the importance of G4s for transcriptional and translational regulation that has previously been recognized (Huppert & Balasubramanian 2007), (Arora et al. 2008). We provide several lines of evidence, including a high enrichment at splice site regions, preference for the non-template strand and *in vitro* experiments, to suggest that G4s form at the pre-mRNA and can modulate alternative splicing events. In addition, RNA G4s are more stable than DNA G4s, suggesting that they could have a greater influence in the transcriptome than in the genome (Arora et al. 2008), (Joachimi et al. 2009). However, co-transcriptional splicing has been previously demonstrated to be the norm (Neugebauer 2002), (Shukla & Oberdoerffer 2012), and we cannot rule out the possibility that the nascent transcripts form DNA-RNA hybrids, implying more complex interactions (Xiao et al. 2013).

Weaker splice sites lead to suboptimal exon recognition, which enable alternative splicing to be modulated by additional cis-regulatory elements or epigenetic factors (Ast 2004). Here we show that there is a pronounced enrichment of G4s at weak splice sites and provide evidence for widespread contribution of G4 structures over alternative splicing. We also find that G4s appear in a subset of species near splice sites (Fig 6) which suggests that they have emerged as splicing modulators during vertebrate evolution. The presence of additional regulatory mechanisms is in accordance with higher frequencies of alternative splicing events in vertebrates compared to invertebrates (Artamonova & Gelfand 2007). Moreover, G4s display a higher likelihood of DNA mutations (Du et al. 2014) and as a result they are likely plastic in nature, enabling rapid splicing changes during evolution and the establishment of new functions through alternative splicing and the generation of isoform diversity.

We observed widespread exon skipping following potassium depolarisation of neurons (Fig 5a, S8), a phenomenon that to our knowledge has only been documented for a handful of cases (An & Grabowski 2007), (Lee et al. 2007), (Liu et al. 2012), (Fiszbein & Kornblihtt 2017). These alternative splicing changes are likely induced by Ca^2+^ influx after depolarisation that is known to affect splicing via CaMK IV (Sharma & Lou 2011). Here, we show that the changes in splicing patterns are associated with the presence of G4s at the splice junctions. However, it is unclear by which mechanism G4s influence alternative splicing. One plausible explanation is that the folded G4 masks otherwise accessible RBP sites. In further support of this possibility, it has been shown that increased unwinding activity of RNA helicases results in fewer G4s being formed (Decorsière et al. 2011), (Haeusler et al. 2014), (Huang et al. 2017). Therefore, inclusion of exons flanked by G4s could involve the interplay between the linear form of the G4 motif and splicing factors, such as hnRNP H/F (Decorsière et al. 2011), (Huang et al. 2017). While the secondary structure formation could conceal signals associated with hnRNP H/F binding (Samatanga et al. 2013), other experimental evidence suggests that G4 stabilisation promotes hRNP F binding (Huang et al. 2017). We hypothesize that the splicing outcome is determined by competition between the kinetics of RBP-RNA interaction and G4 formation (Zhang et al. 2019). A similar type of mechanism has been reported for transcriptional regulation, an example being the promoter G4 at *CMYC* acting as a molecular switch (Siddiqui-Jain et al. 2002). As evidenced by the analysis of KCl RNA-seq experiments, we propose that G4s can function as splicing switches, reinforcing the notion that G4s act as systemic response elements to cellular changes, such as ionic concentrations alterations.

The fact that G4 formation at the RNA level can be modulated by helicases, monovalent ions or small molecules (Balasubramanian & Neidle 2009), (Neidle 2016), (Tippana et al. 2019), (Zhang et al. 2019) opens up new avenues for modulating splicing for therapeutic purposes. It is plausible that by perturbing the stability of G4s or the lifetime of the folded state other RBP binding sites can either become exposed or masked, modulating alternative splicing events. Drugs targeting splicing modulation are already clinically approved, an example being Spinraza for spinal muscular atrophy (Meijboom et al. 2017). There are already multiple compounds available with varying specificities for G4 binding. One example is Quarfloxin, which previously reached phase II clinical trials targeting a G4 in the *CMYC* promoter, but its evaluation was halted due to interference with pol I in rRNA. (Drygin et al. 2009). Our work suggests that G4s in splice sites could be used as a novel pharmacological strategy.

## Acknowledgements

We thank Christopher W.J. Smith and Roberto Munita for important discussions and critical feedback on the manuscript. IGS, GEP and MH are supported by the Wellcome Sanger Institute core grant. HYW and CKK are supported by Research Grants Council of the Hong Kong SAR, China Project No. CityU 11100218, N_CityU110/17, and CityU 21302317, Croucher Foundation Project No. 9500030. Additionally, this work was supported by Cancer Research UK (C13474/A18583, C6946/A14492 to EAM) and the Wellcome Trust (104640/Z/14/Z, 092096/Z/10/Z to EAM).

## Author contributions

IGS and GEP conceived the study and carried out the computational analysis, supervised by EAM and MH. HYW carried out the validation experiments, supervised by CKK. IGS, GEP and MH led the writing of the manuscript with input from HYW, CKK and EAM.

## Supplementary Material

**Supplementary Figure 1:**
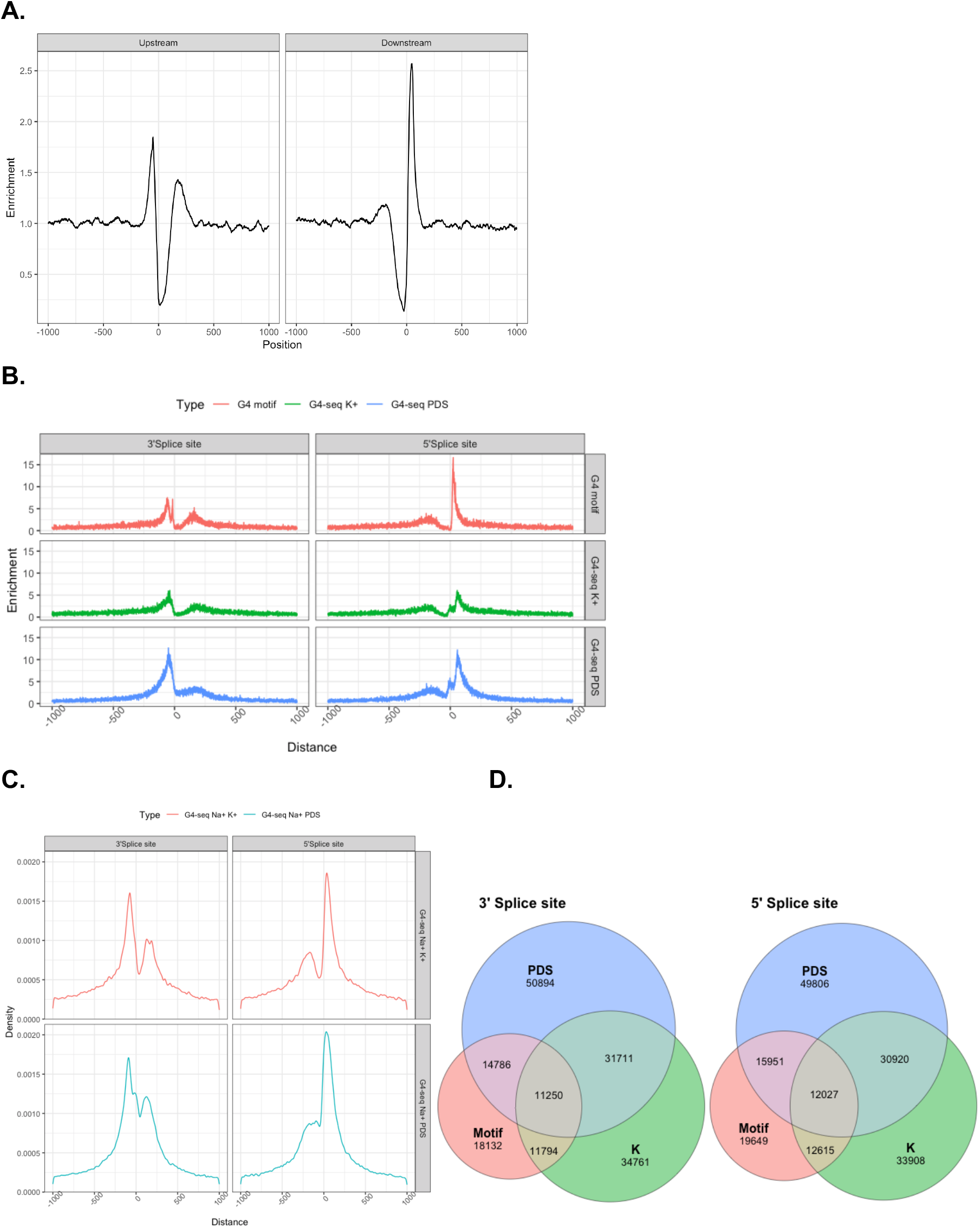
Distribution of G4s in the vicinity of splice sites. **A.** Distribution of G4 motifs at 5’ / 3’ splice sites in mouse. **B.** Enrichment between nearest G4 motif / G4-seq derived G4 peak and a splice site separately for 3’ss (upstream) and 5’ss (downstream) using G4-seq datasets from (Marsico et al. 2019). **C.** Distribution of G4-seq derived G4 peaks within 1kB of the 3’ss and 5’ss after treatment with Na^+^-PDS and Na^+^-K^+^. **D.** Venn diagrams for the number of G4 motifs found in a 100 nt window at the 3’ss and 5’ss that overlap G4-seq derived G4 peaks after treatment with Na^+^-K^+^ and Na^+^-PDS. G4-seq data with Na^+^-PDS and Na^+^-K^+^ treatments were taken from (Chambers et al. 2015).

**Supplementary Figure 2:**
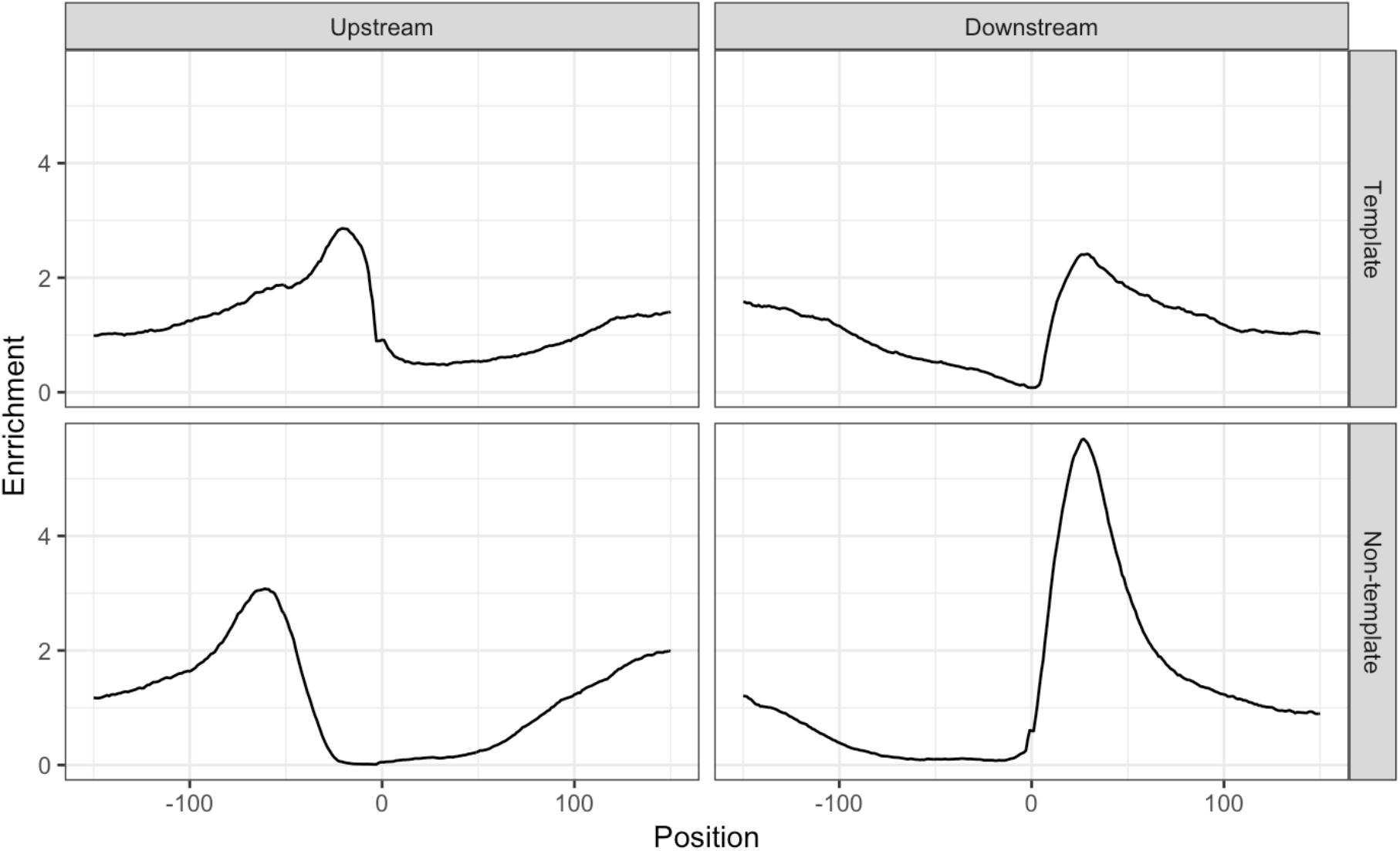
The distribution of G4s at splice junctions differs between the template and non-template strands. The distribution of G4s at splice junctions differs between the template and non-template strands.Distribution of G4 motifs at template and non-template strands at 5’ss (upstream) and 3’ss (downstream).

**Supplementary Figure 3:**
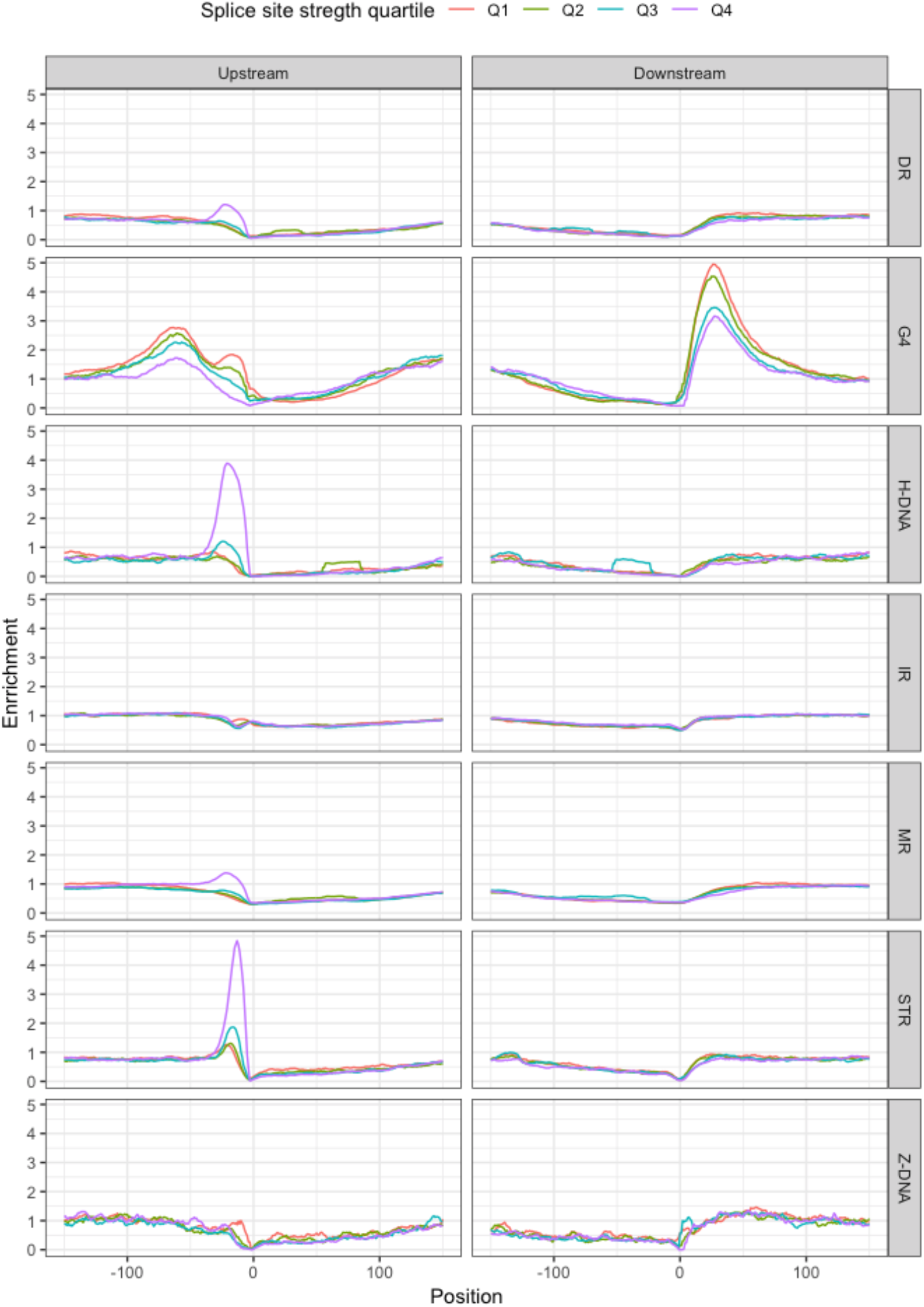
Splice site strength and distribution of non-B DNA motifs at splicing sites. Dependence upon the splicing strength for seven non-B DNA motifs and their distribution at splicing sites relative to the 3’ss (upstream) and the 5’ss (downstream).

**Supplementary Figure 4:**
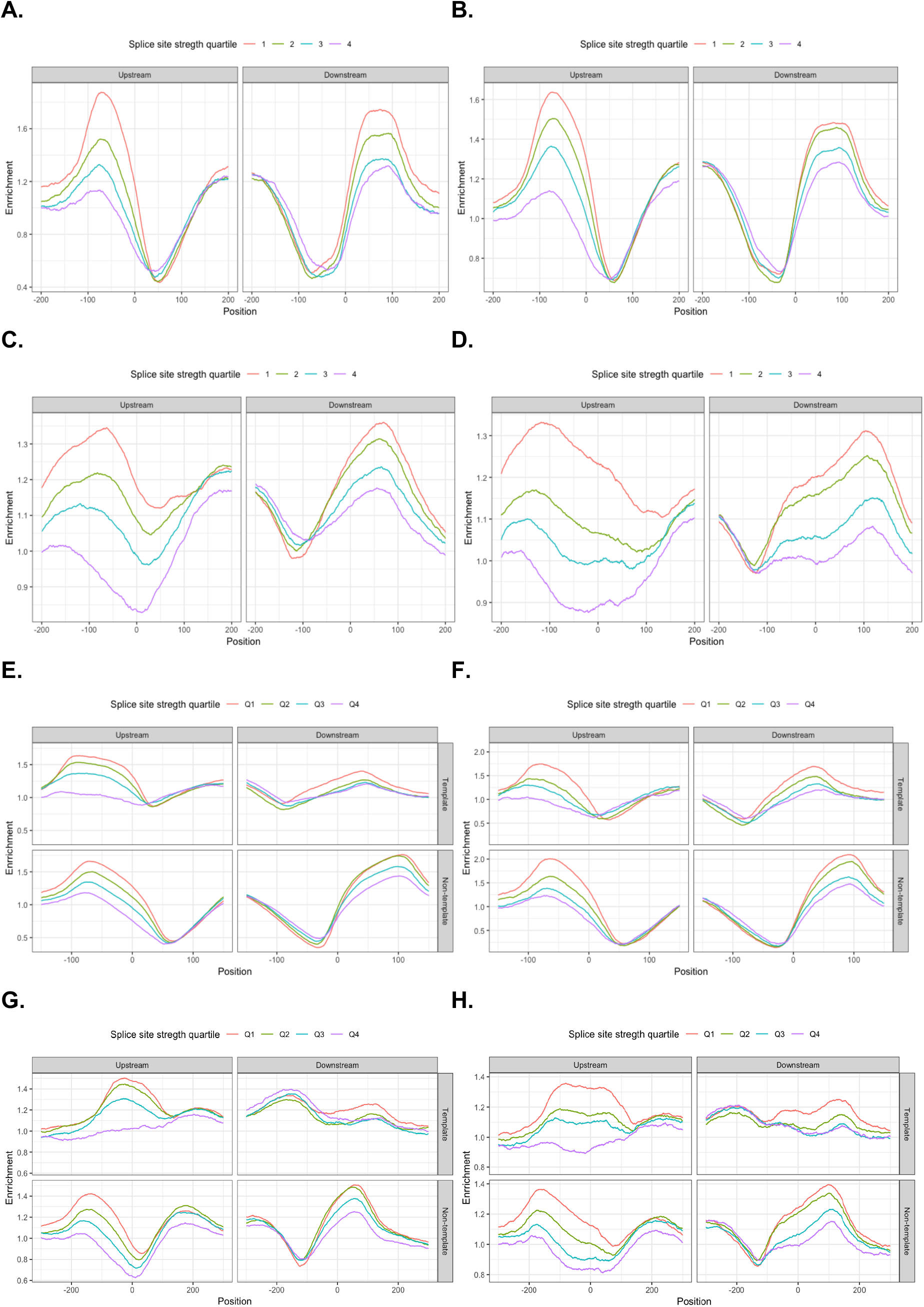
Splice site strength and distribution of G4 motifs at splicing sites. G4s display a stronger enrichment at weaker splice sites. **A.** Distribution of G4 peaks derived from G4-seq with K^+^ treatment from (Marsico et al. 2019) at splicing sites at 3’ss (upstream) and 5’ss (downstream) and the association with splicing strength. **B.** Distribution of G4 peaks derived from G4-seq with PDS treatment from (Marsico et al. 2019) at 3’ss (upstream) and 5’ss (downstream) and the association with splicing strength. **C.** Distribution of G4 peaks derived from G4-seq with Na^+^-K^+^ treatment from (Chambers et al. 2015) at 3’ss (upstream) and 5’ss (downstream) and the association with splicing strength. **D.** Distribution of G4 peaks derived from G4-seq with Na^+^-K^+^ treatment from (Chambers et al. 2015) at 3’ss (upstream) and 5’ss (downstream) and the association with splicing strength. **E.** Distribution of G4 peaks derived from G4-seq in presence of PDS at template and nontemplate strands and the association with splicing strength. **F.** Distribution of G4 peaks derived from G4-seq in presence of K^+^ at template and non-template strands and the association with splicing strength. **G.** Distribution of G4 peaks derived from G4-seq in presence of Na^+^-PDS at template and non-template strands and the association with splicing strength. **H.** Distribution of G4 peaks derived from G4-seq in presence of Na^+^-K^+^ at template and non-template strands and the association with splicing strength. A-B and E-F panels used recent G4-seq data from (Marsico et al. 2019) with higher resolution than C-D and G-H panels which used older G4-seq data from (Chambers et al. 2015). Median peak length for G4-seq peaks from (Chambers et al. 2015) was 255 nt and 195 nt in Na^+^-K^+^ and Na^+^-PDS conditions, respectively. Median peak length for G4-seq peaks from (Marsico et al. 2019) was 135 nt and 120 nt in PDS and K^+^ conditions, respectively.

**Supplementary Figure 5:**
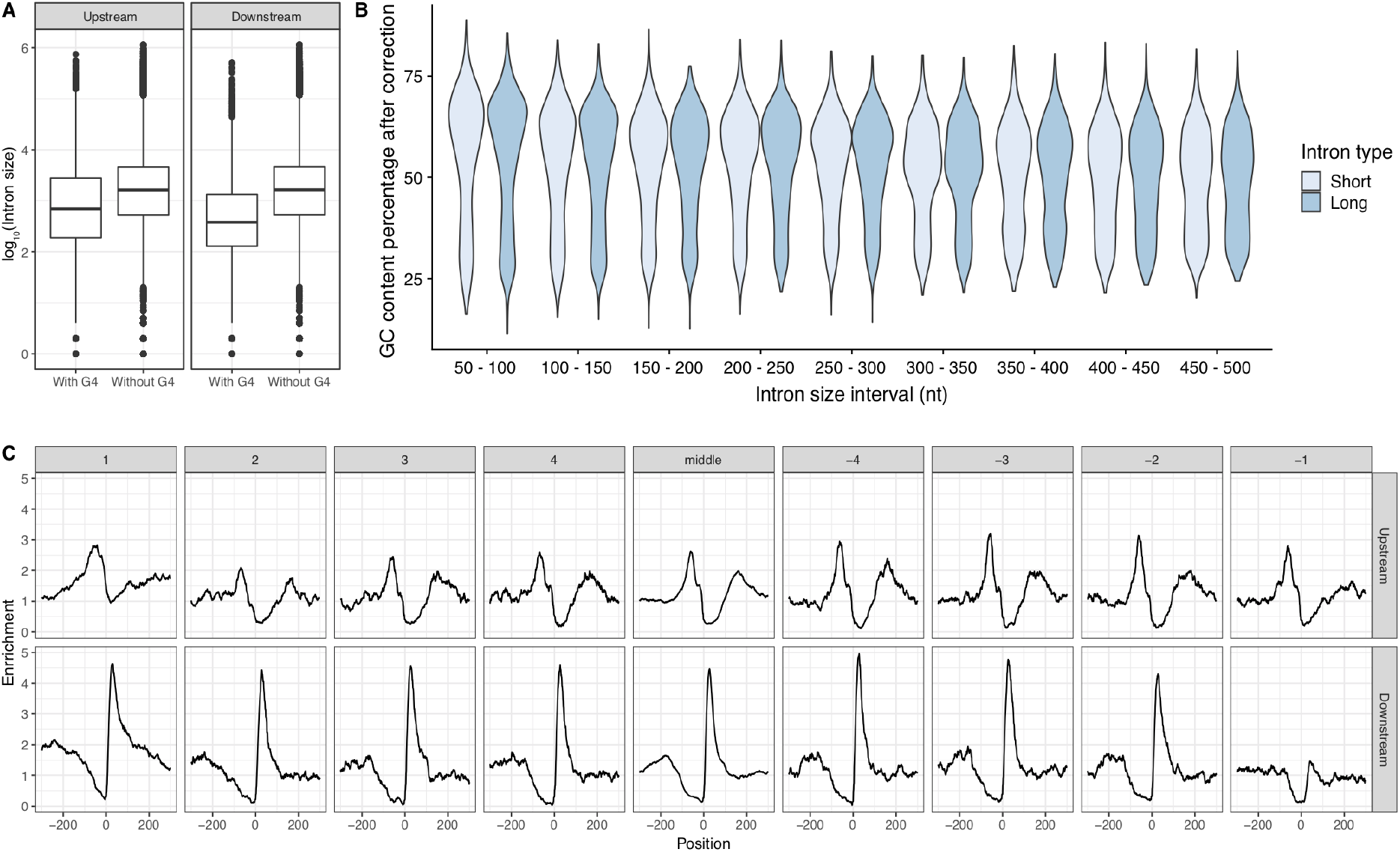
**A.** Intron size of the upstream and downstream introns was calculated for groups with or without a G4 within 100 bps of the splice site (Mann-Whitney U p-value <0.001 for both upstream and downstream introns). **B.** GC content distribution across selected groups of short and long introns. Intron size interval refers to the size of small introns. Long introns were defined as introns > 500 bp. **C**. G4 motif enrichment relative to the splice site across exons in the gene body at the 3’ss (upstream) and at the 5’ss (downstream). G4 motifs are enriched at both 3’ss and 5’ss across splice sites throughout the gene body. Exons were separated intro first to fourth exons, middle exons, last four exons and the distribution of G4s was studied individually at each of them.

**Supplementary Figure 6:**
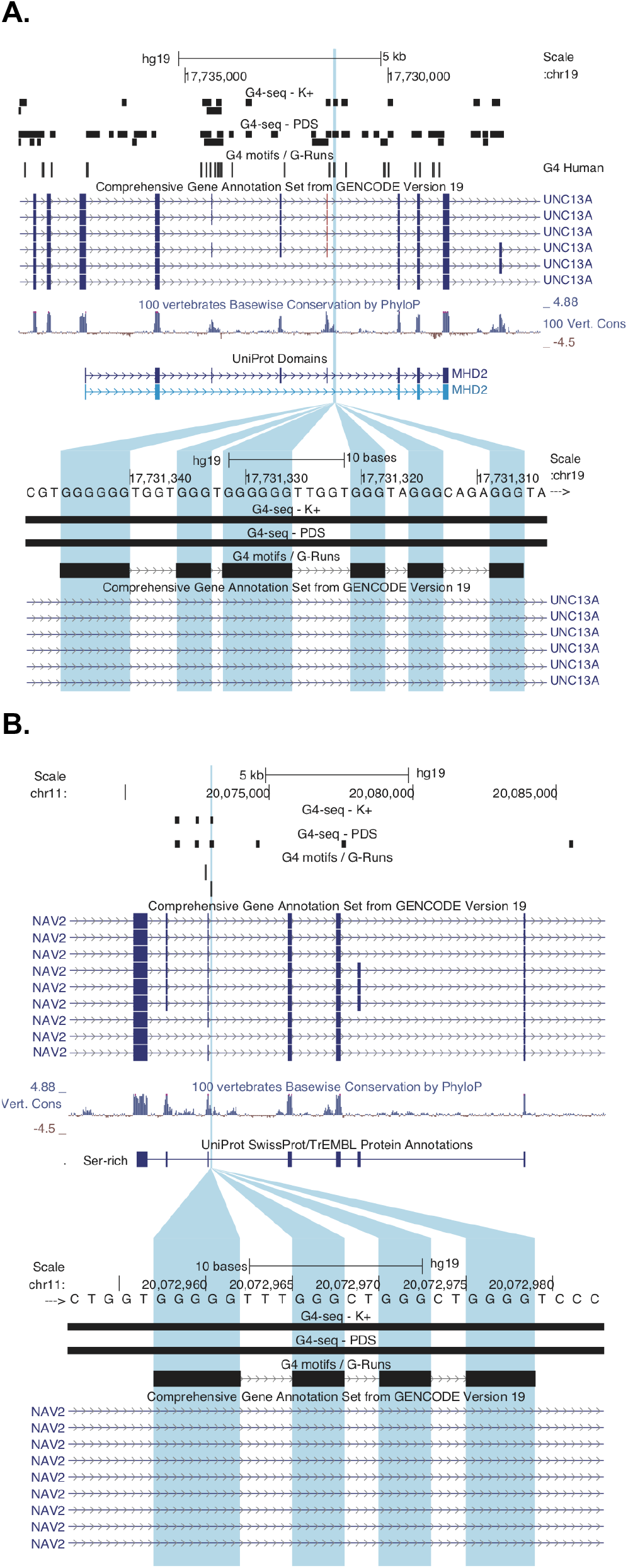
Illustration of two loci with exons flanked by G4s, displaying alternative splicing changes following KCl treatment. **A.** Schematic of *UNC13A* exon 38 and the G4s around its splice junction. **B.** Schematic of *NAV2* exon 16 and the G4s around its splice junction.

**Supplementary Figure 7:**
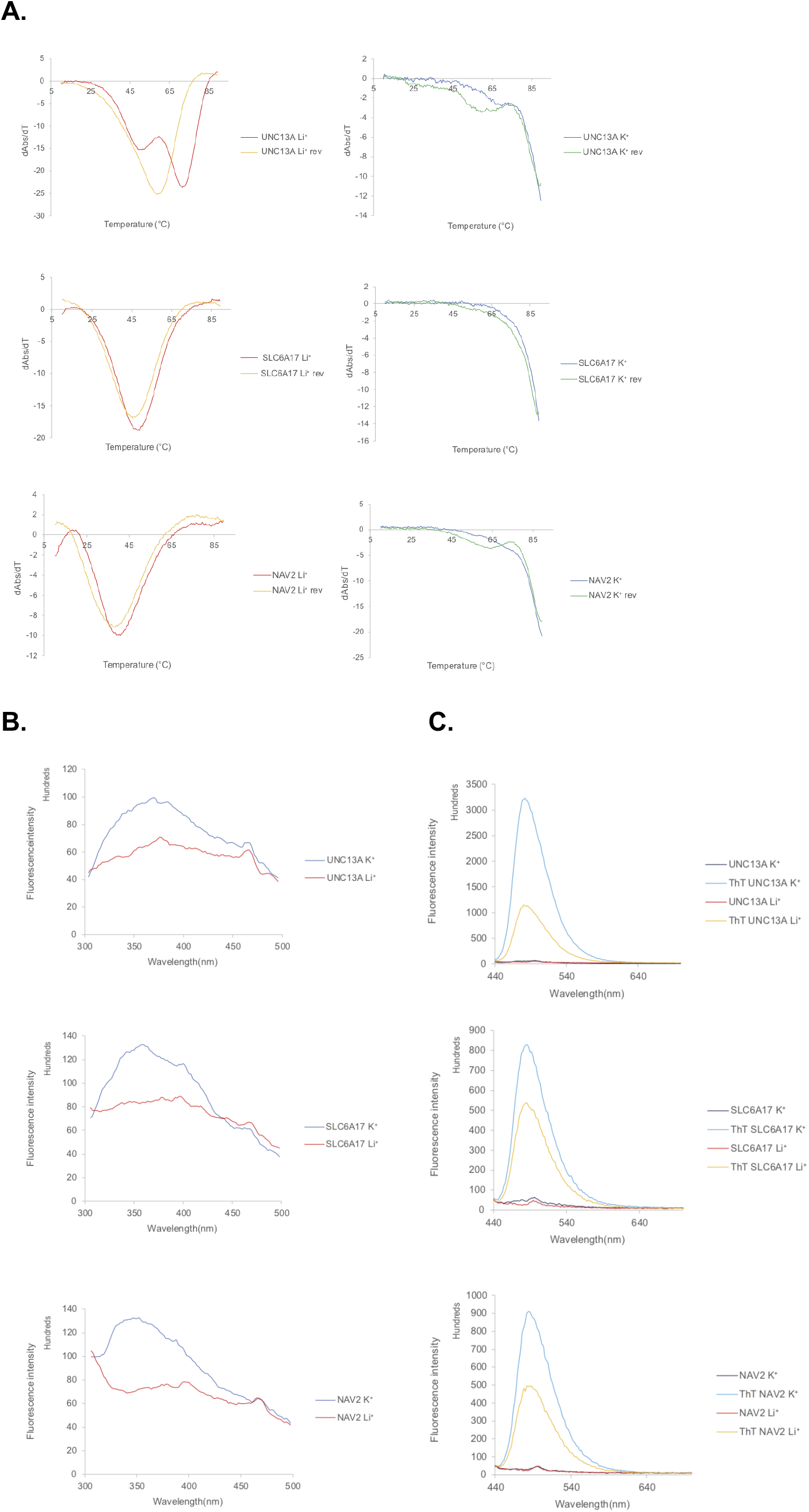
G-quadruplex structure formation identification through fluorescence emission in presence of G4 stabilising (K^+^) and non-stabilising (Li^+^) cations. **A.** UV melting profiles of the three G4 candidates in presence of Li^+^ and K^+^. Both the forward and reverse melt were shown here **B.** Intrinsic fluorescence of the three candidate RNA oligonucleotides under Li^+^ or K^+^ conditions. The intrinsic fluorescence of G4s was increased when replacing Li^+^ with K^+^, highlighting the formation of RNA G4s. **C.** ThT ligand enhanced fluorescence of the three candidate RNA oligonucleotides under Li^+^ or K^+^ condition. In the absence of ThT ligand, no fluorescence was observed at ~480nm. Upon ThT addition, weak fluorescence was observed under Li^+^, which was substantially enhanced when substituted with K^+^, supporting the formation of G4 which allow recognition of ThT and enhance its fluorescence.

**Supplementary Figure 8:**
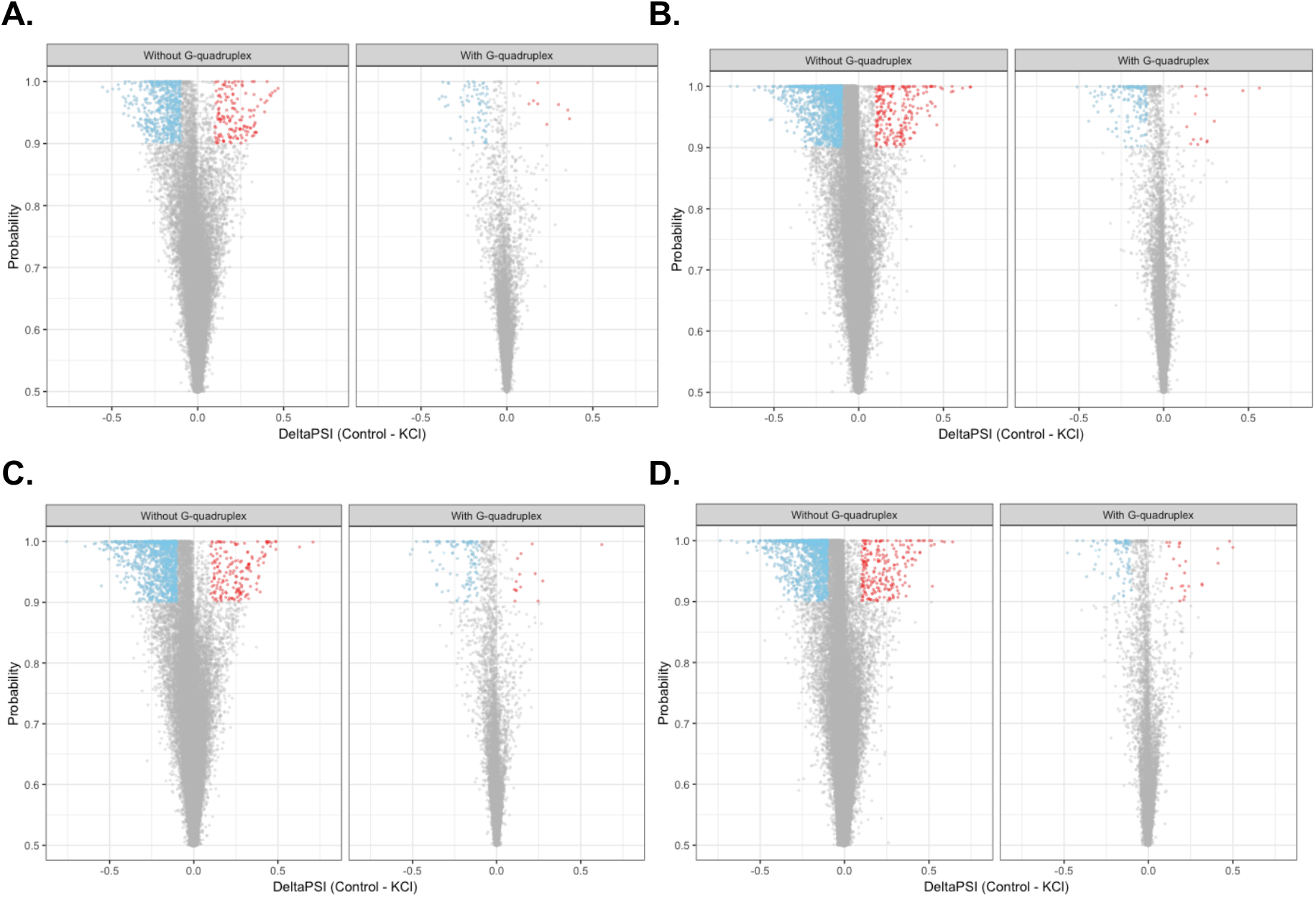
Neuronal stimulation with KCl results in exon skipping in human and mouse neuronal cells. Volcano plots for **A.** CD1 mouse ESC-derived neurons, **B.** CD1 mouse primary cortical neurons at DIV4, **C.** CD1 mouse primary cortical neurons at DIV10, **D.** Tc1 mouse primary cortical neurons at DIV10. We found consistent exon skipping of canonical exons following KCl treatment in all cases (binomial tests with Bonferroni correction: p-value<0.001 in all cases).

**Supplementary Figure 9:**
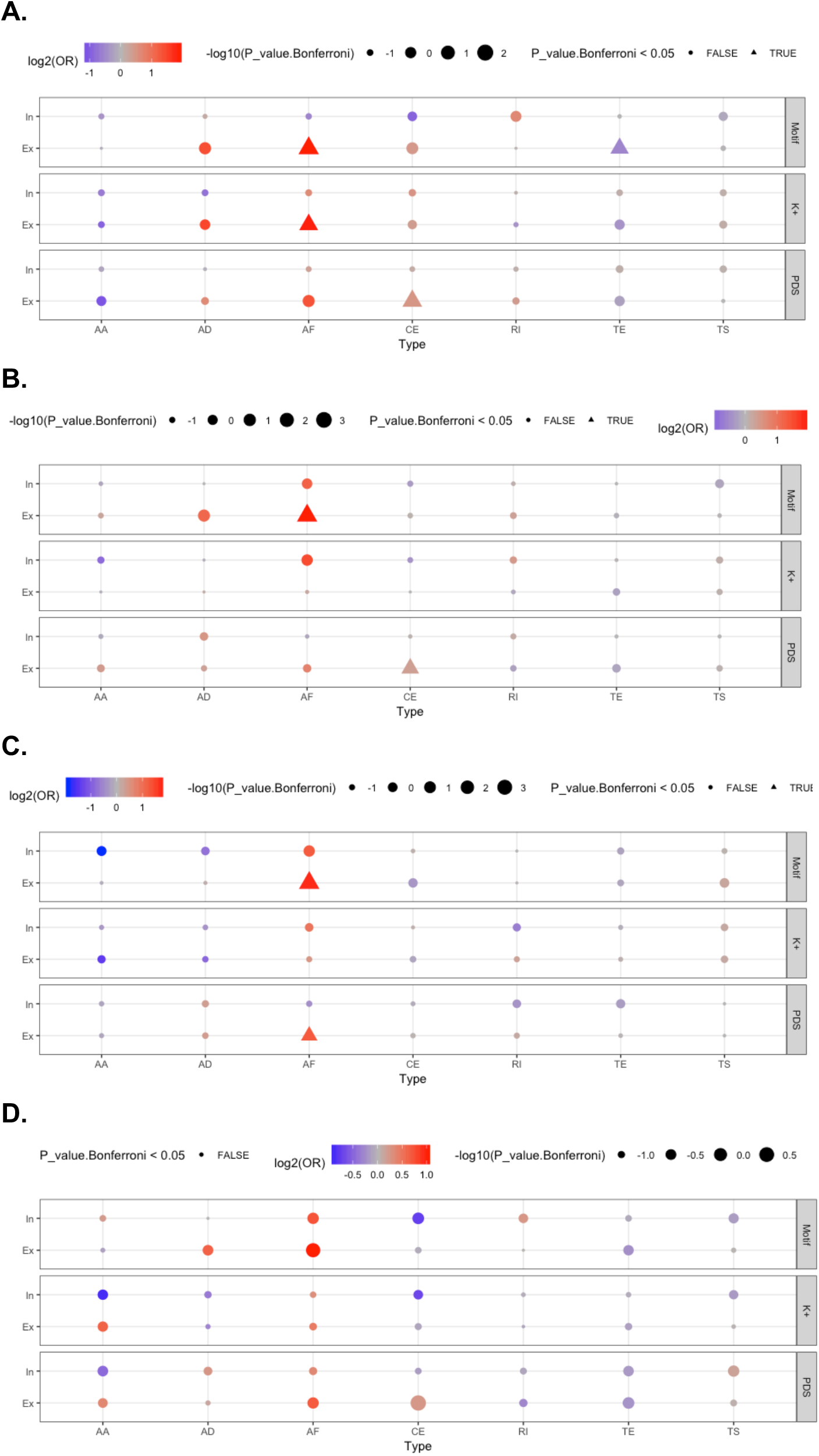
Inclusion and exclusion patterns of genic regions in association to G4 presence or absence following KCl treatment. All p-values were generated with Chi-squared test and were Bonferroni corrected. **A.** ESC-derived neurons from CD1 mice. **B.** Primary cortical neurons from DIV10 CD1 mice. **C.** Primary cortical neurons from DIV10 Tc1 mice. **D.** Primary cortical neurons from DIV4 CD1 mice. Types correspond to: AA – Alternative Acceptor Splice Site, AD - Alternative Donor Splice Site, AF - Alternative First Exon, CE – Core Exon, TE - Tandem Alternative Polyadenylation Site, TS - Tandem Transcription Start Site.

**Supplementary Figure 10:**
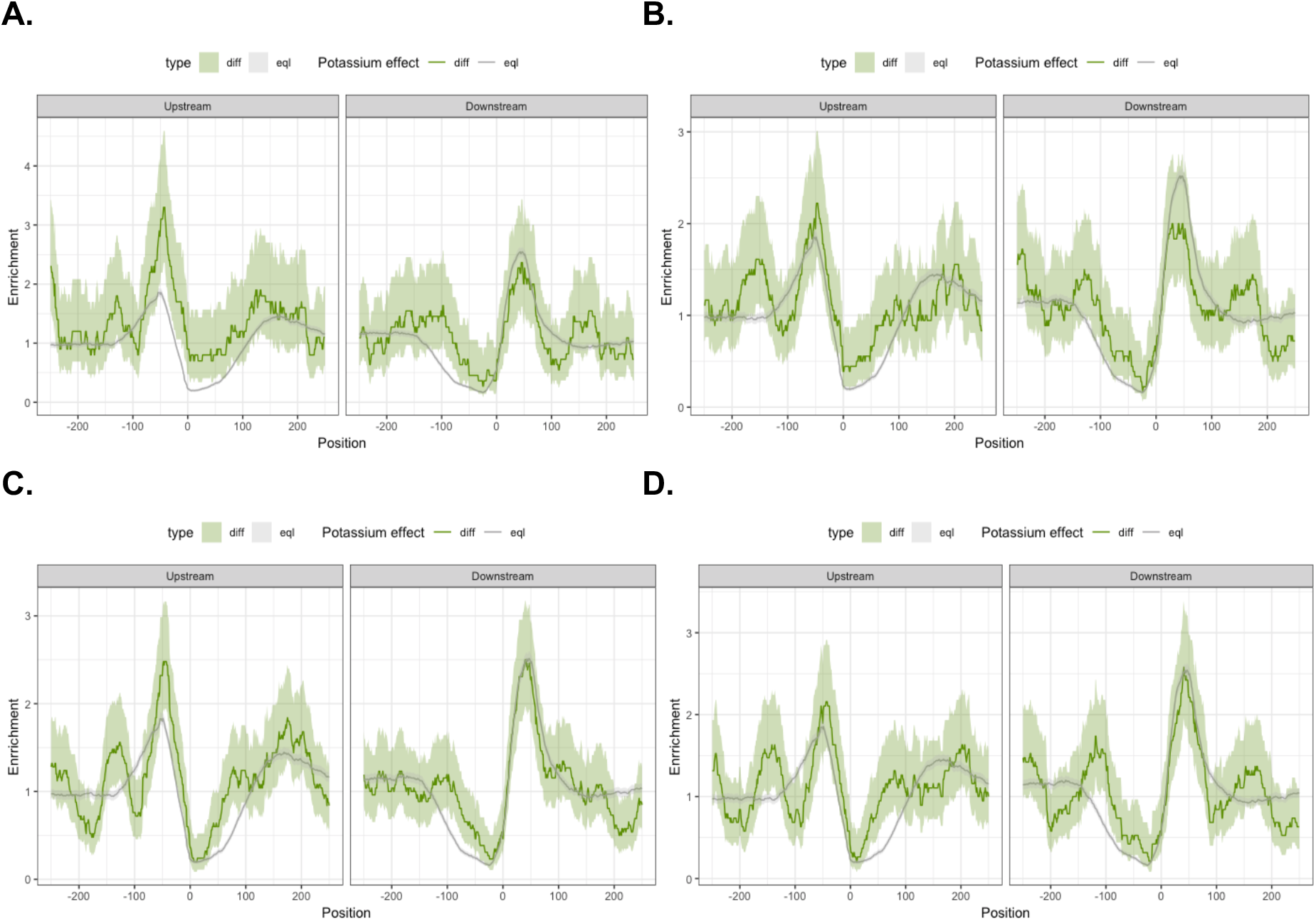
Neuronal stimulation with KCl results in G4-mediated alternative splicing. **A.** Distribution of consensus G4 motifs at splicing sites after potassium depolarisation of CD1 mouse ESC-derived neurons. **B.** Distribution of consensus G4 motifs at splicing sites after potassium depolarisation of Tc1 mouse neurons at DIV10. **C.** Distribution of consensus G4 motifs at splicing sites after potassium depolarisation of CD1 mouse neurons at DIV4. **D.** Distribution of consensus G4 motifs at splicing sites after potassium depolarisation of CD1 mouse neurons at DIV10. The error bands represent 0.95 confidence intervals.

**Supplementary Figure 11:**
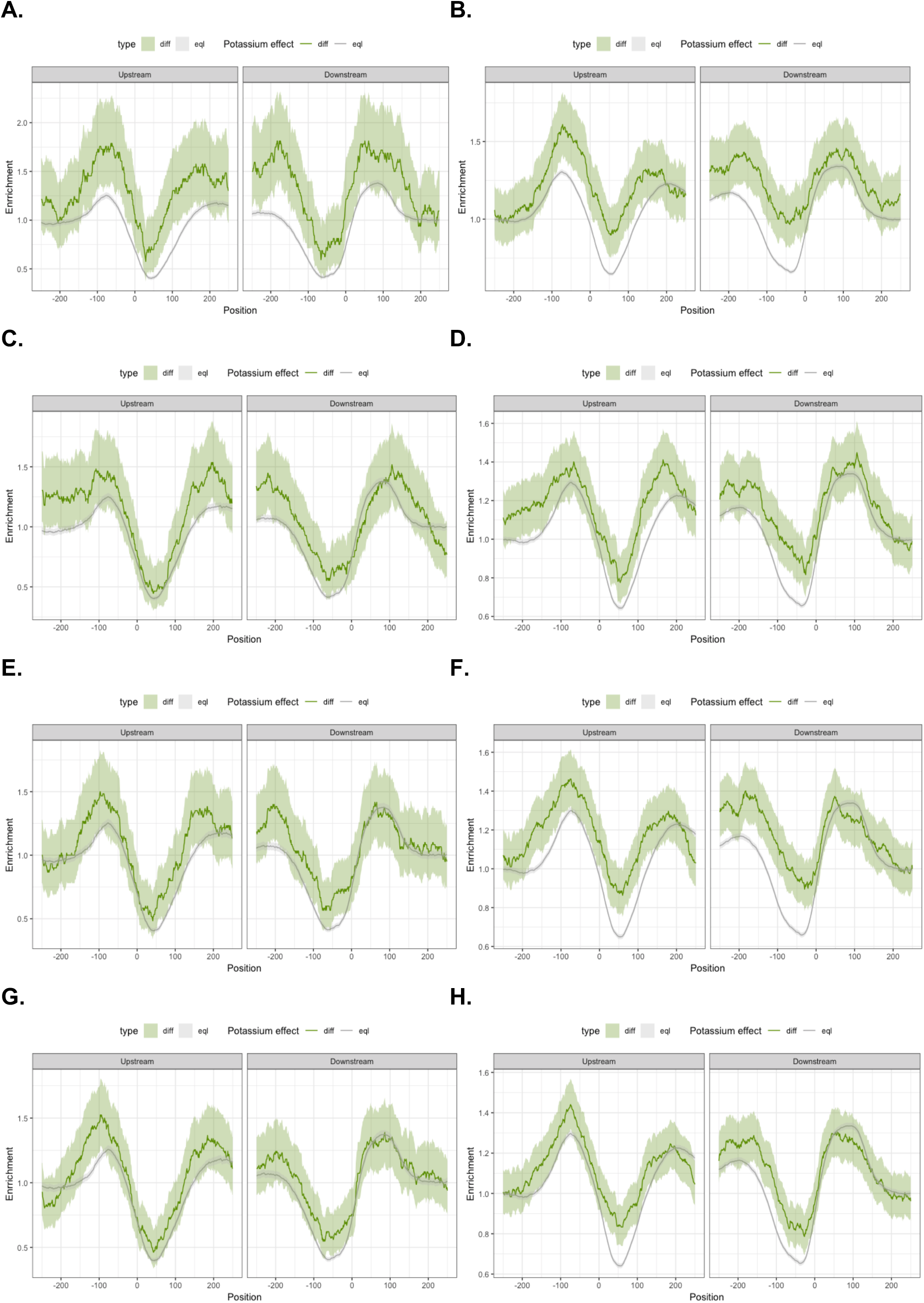
Neuronal stimulation with KCl results in G4-mediated alternative splicing using G4-seq derived G4 peaks. **A.** Distribution of G4-seq derived G4 peaks with K^+^ treatment at splicing sites after potassium depolarisation of CD1 mouse ESC-derived neurons. **B.** Distribution of G4-seq derived G4 peaks with PDS treatment at splicing sites after potassium depolarisation of CD1 mouse ESC-derived neurons. **C.** Distribution of G4-seq derived G4 peaks with K^+^ treatment at splicing sites after potassium depolarisation of Tc1 mouse neurons at DIV10. **D.** Distribution of G4-seq derived G4 peaks with PDS treatment at splicing sites after potassium depolarisation of Tc1 mouse neurons at DIV10. **E.** Distribution of G4-seq derived G4 peaks with K^+^ treatment at splicing sites after potassium depolarisation of CD11 mouse neurons at DIV10. **F.** Distribution of G4-seq derived G4 peaks with PDS treatment at splicing sites after potassium depolarisation of CD1 mouse neurons at DIV10. **G.** Distribution of G4-seq derived G4 peaks with K^+^ treatment at splicing sites after potassium depolarisation of CD1 mouse neurons at DIV4. **H.** Distribution of G4-seq derived G4 peaks with PDS treatment at splicing sites after potassium depolarisation of CD1 mouse neurons at DIV4. The error bands represent 0.95 confidence intervals.

**Supplementary Figure 12:**
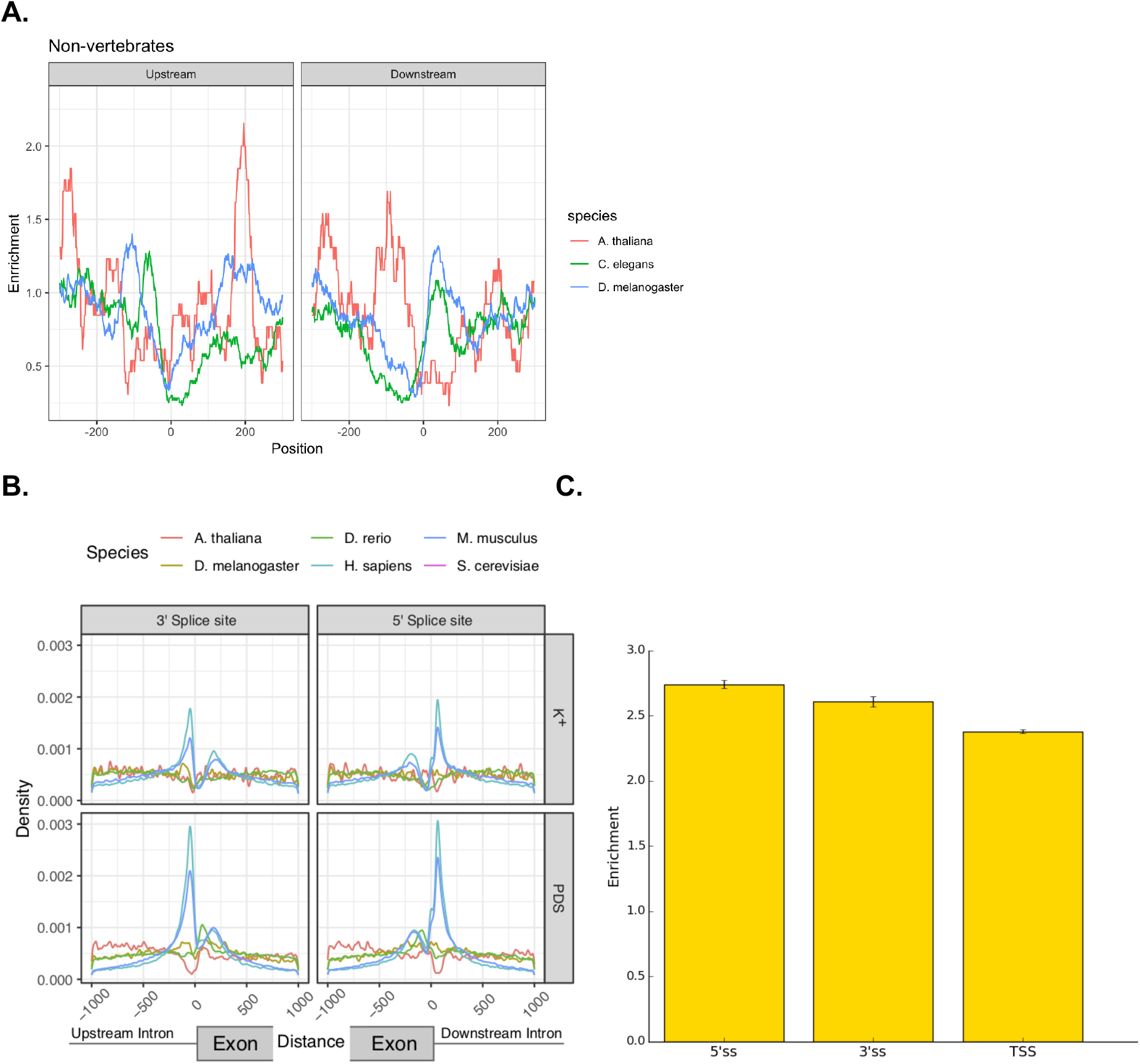
Distribution of G4s at splicing sites across multiple species. **A.** Enrichment of consensus G4 motifs at splicing sites across different non-vertebrate species. Enrichment indicates the frequency of G4s at a position over the median frequency across the window. **B.** Density of G4s at: 5’ss and 3’ss across six species for PDS and K^+^ treatments. **C.** Enrichment (Observed / Expected) of consensus G4 motifs at 3’ss, 5’ss and TSS for humans. Observed frequency was calculated at 100 nt windows each side of the 3’ss, 5’ss and TSS sites and expected frequency was calculated across 1000-fold simulations of all 100 nt windows controlling for dinucleotide content.

## 4. Materials and Methods

### 4.1. Genome and gene annotations processing

We obtained genome assemblies from the UCSC Genome Browser FTP server for eleven organisms: GRCh37 (hg19) reference assembly of the human genome, the mouse reference genome (mm10), the *Saccharomyces cerevisiae* reference genome (sacCer3), the chicken reference genome (galGal5), the *Drosophila melanogaster* reference genome (dm6), the zebrafish reference genome (danRer11), the *Xenopus tropicalis* reference genome (xenTro9), the *Anolis carolinensis* reference genome (anoCar2), the Arabidopsis thaliana reference genome (Tair10) and the *Caenorhabditis elegans* reference genome (ce10).

We downloaded the Ensembl gene annotation files for the associated genomes from UCSC Table Browser as BED files for each species (Karolchik et al. 2003). Using in-house python scripts we extracted the coordinates of internal exons flanked by canonical splice sites (GT-AG introns) for every species. To calculate the splicing strength scores, we used publicly available positional frequency matrices from the SpliceRack database (Sheth et al. 2006) and previously developed scripts used before for the same purpose (Parada et al. 2014). Splice sites were grouped into quartiles based on their splicing strength score for the downstream analyses to study the distribution of non-B DNA motifs and in particular G4 motifs (Fig 2a), (S3), (S4). For figure 2a, the confidence intervals were calculated using “binconf” command from “Hmisc” package in R with default parameters. Mann-Whitney U tests were performed at 100 nt each side in the upstream splice site and at the downstream splice site to compare the splicing strength scores of sites with and without G4s. Phylogenetic trees were generated using PhyloT software (Fig 4a), (S6a-b).

### 4.2. Genomic datasets

#### Non-B DNA motifs

Identification of each non-B DNA motif was performed using the genomewide maps in humans and mice provided by (Cer et al. 2013) and processed as described in (Georgakopoulos-Soares et al. 2018). We focused on seven non-B DNA motifs; inverted repeats, mirror repeats, H-DNA which forms at a subset of mirror repeats with high AG content, G4s, Z-DNA which forms at non-AT alternating purine pyrimidine stretches, short tandem repeats and direct repeats (Fig 1a), (S3).

Regular expressions were employed to identify genome-wide consecutive G-runs across the human genome, interspersed with loops of up to 7 bps. In total, one to six consecutive G-runs were searched (Fig 2b). For each species we generated the genome-wide G4 maps using a regular expression of the consensus G4 motif (G⩾3Nl-7G⩾3Nl-7G⩾3Nl-7G⩾3), (Fig 6a-b). Orientation of G4s and G-runs was performed with respect to template and non-template strands to calculate strand asymmetries at genic regions as previously described for polyN motifs (N being Gs, Cs, Ts and As) in (Georgakopoulos-Soares et al. n.d.),(Fig 2a), (S2), (S4e-h).

Permuted windows of 100 nt each side of each splice junction were generated using ushuffle (Jiang et al. 2008) correcting for dinucleotide content. The fold enrichment for G4s was calculated as the ratio of the number of motifs found in the real sequences and the median of 1,000 permutations of the set of all real sequences (S12c). The corrected enrichment of G4s at 3’ss and 5’ss was calculated as the ratio of the real enrichment of G4s over the background enrichment of G4s at shuffled splice site windows.

To investigate the relationship between non-B DNA motifs or G4-seq peaks and splice sites we generated local windows around the splice sites and measured the distribution of each non-B DNA motif (Fig 1a), (S3) or G4-seq dataset across the window (Fig 1b), (S1a,c). The enrichment was calculated as the number of occurrences at a position over the median number of occurrences across the window. Regardless of the window size shown in figures, the enrichment was calculated over a window of 1kB. The same approach was used to calculate the enrichment of G4s at splice sites across different species (Fig 6b).

The density of G4 consensus motifs or G4-seq derived peaks at local windows was calculated as the number of occurrences of the motif or the peak over the total number of base pairs examined (Fig 6a,c).

#### G4-seq data

G4-seq BedGraph data were obtained from GEO accession code GSE63874 (Chambers et al. 2015) for the human genome and analyzed with bedtools closest command to identify the closest G4 to splice sites and to calculate the distance. The analysis was performed separately for Na^+^-K^+^ and Na^+^-PDS conditions and it was compared to the distribution obtained from the G4 consensus motif. G4-seq BedGraph data for six species, human, mouse, *D. melanogaster, C. elegans, A. thaliana* and yeast, were obtained from GEO accession code GSE110582 (Marsico et al. 2019) and analyzed using the same genome annotations as those used for the generation of each G4-seq dataset.

Coordinates for internal exons flanked by canonical splice sites (GT-AG introns) were extracted for each species using the Ensembl annotation versions described in (Marsico et al. 2019) using custom python scripts.

#### Relationship between G4s and exon / intron length

Introns and exons were grouped based on the presence or absence of G4s within 100 nt each side of the 5’ss and 3’ss and further subdivided into those containing a G4 on the template or on the non-template strand, separately for the 3’ss and the 5’ss. For each of the eight groups we calculated the median length of the intron or exon in a group and performed Mann-Whitney U tests to calculate the significance of the association between length of exons / introns and G4 presence (Fig S5a). The R function stat_density was used to plot the length distribution of introns with and without G4s as modelled by a kernel density estimate (Fig 3a). Abundance enrichment of intron length in 3’ / 5’ splice sites in relationship with presence of G4s was generated in R using the function geom_smooth in an eighth grade model (Fig 3b). Correction of GC content in introns with different length was performed by grouping introns into small introns (<500nt) and large introns (>500nt). Then we calculated the GC content for both groups and for each short intron we selected a long intron with a close GC content value, in such a way that GC distribution across short and long introns groups were nearly identical (S5b).

#### G4s and relationship to exon number

For the longest transcript of each gene with nine or more exons we separated exons into 9 groups, the first four exons, the last four exons and the remaining middle exons. To compare the frequency of G4s in splice junctions across the gene body we calculated the distribution of G4s in each exon group relative to the 5’ / 3’ splice sites (S5c). We also calculated the distribution of G4s in each exon group relative to the 5’ / 3’ splice sites separately for the template and non-template strands (Fig 3e).

#### Relationship between G4s, splicing strength score and intron length

We calculated the splicing strength score and intron length for the upstream and downstream intron of each exon. We separated introns and splicing strength scores into deciles and calculated the G4 density at each decile, from which we produced two heatmaps displaying the density of G4s as a function of splicing strength score and intron length for the upstream and downstream introns (Fig 3c).

### 4.3. Comparative analysis of RNA-seq experiment

#### Differential exon inclusion following depolarisation

We analyzed available data (BioProject Accession: PRJEB19451, ENA link: ERP021488) for mouse and human ESC-derived cortical neurons, mouse primary cortical neurons from wild-type and Tc1 mice stimulated with KCl treatment and untreated followed by RNA-seq four hours post-treatment (Qiu et al. 2016). We used Whippet (Sterne-Weiler et al. 2018) to quantify splicing nodes and asses their alternative inclusion after KCl treatment and controls (Fig 5a-b), (S8-9). We used absolute value of DeltaPSI greater than 0.1 and probability greater than 0.9 to define a splicing node as differentially included between treatments and controls.

We calculated the distance between the middle point of G4 motifs and G4-seq peaks from each splicing node to determine their association with G4s. Splicing nodes whose splice sites were within 100 bps to G4 motif or 45 bps to G4-seq peak were classified as G4 associated splicing nodes. Next, we assessed the influence of G4s to splicing changes following KCl depolarisation of human and mouse neurons by calculating the odds ratio score of each splicing node type. To determine the statistical significance of the effect we performed chi-squared test using Yates’ correction and also adjusting for multiple testing with Bonferroni corrections (Figure 5a, S9). The distribution of G4 motifs and G4-seq peaks was profiled around differentially included and non-differentially included core exon splicing nodes (CE), (Figure 5d-e, S10–S11). The confidence intervals were calculated using “binconf” command from “Hmisc” package in R with default parameters. Sashimi plots were generated using “ggsashimi” package (Garrido-Martín et al. 2018). Inclusion and exclusion path ratios were calculated using the total amount of spliced reads supporting each splice junction, where inclusion paths were calculated using the average read count for splice junctions flanking each exon side.

Three putative non-template G4s found in proximity to splicing junctions and which were differentially included following depolarisation in human ESC-derived neurons and in at least one condition in mice were selected for validation experiments. These were: i) a G4 downstream of exon 7 for *SLC6A17* (chr1: 110734886-110734906), ii) a G4 downstream of exon 38 in *Unc13a* (chr19:17731307-17731346) and iii) a G4 upstream of exon 16 in *Nav2* (chr11:20072958-20072979) for which RNA oligonucleotides at the G4 locations were ordered.

The RNA oligonucleotides used were (G-runs marked in bold):

1. SLC6A17 oligonucleotide: **GGG**AGT**GGG**CA**GGGG**T**GGGGG**
2. UNC13A oligonucleotide: **GGGGGG**TGGT**GGG**T**GGGGGG**TTGGT**GGG**TA**GGG**CAGA**GGG**
3. Nrxn2 oligonucleotide: **GGGGG**TTT**GGG**CT**GGG**CT**GGGG**

### 4.4. Experimental validation of G4 candidates

#### NMM ligand enhanced fluorescence

This assay was performed similar to our previous work (Chan et al. 2018). Briefly, sample solutions containing 1 μM RNA were prepared in 150 mM LiCl/KCl, 10 mM LiCac buffer (pH 7.0) and 1 μM NMM ligand. Fluorescence spectroscopy was performed using HORBIA FluoroMax-4 and a 1-cm path length quartz cuvette (Wuxi Jinghe Optical Instrument Co.) was used with a sample volume of 100 μL. Before the measurement, the samples (ligand not added) were denatured at 95 °C for 3 minutes and allowed to cool down at room temperature for 15 minutes. The samples were excited at 394 nm and the emission spectra were acquired from 550 to 750 nm. Data were collected every 2 nm at 25 °C with 5 nm entrance and exit slit widths. Raw ligand enhanced fluorescence spectra were first blanked with the corresponding sample spectra that resembled all chemical conditions except with the absence of the ligand. All calculations mentioned were performed in Microsoft Excel.

#### ThT ligand enhanced fluorescence

This assay was performed similar to our previous work (Chan et al. 2018). Briefly, sample solutions containing 1 μM RNA were prepared in 150 mM LiCl/KCl, 10 mM LiCac buffer (pH 7.0) and 1 μM ThT ligand. Fluorescence spectroscopy was performed using HORBIA FluoroMax-4 and a 1-cm path length quartz cuvette (Wuxi Jinghe Optical Instrument Co.) was used with a sample volume of 100 μL. Before the measurement, the samples (ligand not added) were denatured at 95 °C for 3 minutes and allowed to cool down at room temperature for 15 minutes. The samples were excited at 425 nm and the emission spectra were acquired from 440 to 700 nm. Data were collected every 2 nm at 25 °C with 5 nm entrance and exit slit widths. Raw ligand enhanced fluorescence spectra were first blanked with the corresponding sample spectra that resembled all chemical conditions except with the absence of the ligand. All calculations mentioned were performed in Microsoft Excel.

#### Circular Dichroism (CD) spectroscopy

This assay was performed similar to our previous work (Chan et al. 2019). Briefly, the CD spectroscopy was performed using Jasco J-1500 CD spectrophotometer and a 1-cm path length quartz cuvette (Hellma Analytics) was employed in a volume of 2 mL. Samples with 5 μM RNA (final concentration) were prepared in 10 mM LiCac (pH 7.0) and 150 mM KCl/LiCl. Each of the RNA samples were then thoroughly mixed and denatured by heating at 95 °C for 5 minutes and cooled to room temperature for 15 minutes for renaturation. The RNA samples were excited and scanned from 220 – 310 nm at 25 °C and spectra were acquired every 1 nm. All spectra reported were average of 2 scans with a response time of 0.5 s/nm. They were then normalized to molar residue ellipticity and smoothed over 5 nm. All data was analyzed with Spectra Manager™ Suite (Jasco Software).

#### Thermal melting monitored by UV spectroscopy

This assay was performed similar to our previous work (Chan et al. 2019). Briefly, samples were prepared to a concentration of 10 mM LiCac buffer, 150 mM salt (KCl/LiCl) and 5 μM RNA, with a total volume of 2 mL. Each of the samples was mixed thoroughly and heated at 95 °C for 5 minutes so as to denature the RNA. It was then cooled for 15 minutes at room temperature for renaturation. All UV melting experiments were performed on an Agilent Cary 100 UV-Vis Spectrophotometer, using 1-cm path length quartz cuvette. Before the experiment started, the sample block was first flushed with dry N2 gas and cooled down to 5 °C for 5 minutes. After the sample solutions were loaded to the cuvettes, they were sealed with 3 layers of Teflon^®^ tape to prevent vaporization at high temperature. The samples were scanned from 5 to 95 °C with a temperature incremental rate of 0.5 °C/min. The temperature was hold at 95 °C for 5-minutes before a reversed scan was performed, scanning from 95 to 5 °C with a rate of 0.5 °C/min. The unfolding and folding transitions in both scans were monitored at 295 nm.

Raw data obtained were subtracted by the blank solutions, which contains the same concentrations of LiCac buffer (pH 7.0) and corresponding salt only. It was then smoothed over 11 nm and its first derivative was plotted in Microsoft Excel. The final melting temperature was obtained by averaging the melting temperatures in the forward and reversed scans.

#### Intrinsic fluorescence spectroscopy

This assay was performed similar to our previous work (Chan et al. 2019). Briefly, intrinsic fluorescence spectroscopy was performed using HORIBA FluoroMax-4 and a 1-cm path length quartz cuvette (Hellma Analytics) was used with a volume of 2 mL. Samples with 5 μM RNA were prepared in 10 mM LiCac (pH 7.0) and 150 mM KCl/LiCl. The samples were then denatured at 95 °C for 5 minutes and cooled to room temperature for 15 minutes for renaturation. For the measurement of intrinsic fluorescence of G-quadruplexes, the samples are excited at 260 nm and the emission spectra were acquired from 300 – 500 nm. Spectra were acquired every 2 nm at 25 °C. The bandwidth of the entrance and exit slits was 5 nm. All data were smoothed over 5 nm. Results here are analyzed using Microsoft Excel.

#### Code and data availability

All the associated code used for the generation of figures and presentation of data throughout the manuscript is deposited in github in the following link: https://github.com/hemberg-lab/Georgakopoulos_Soares_and_Parada_2019.

